# Cancer-associated fibroblasts confer ALK inhibitor resistance in *EML4-ALK*-driven lung cancer via concurrent integrin and MET signaling

**DOI:** 10.1101/2024.08.27.609975

**Authors:** Qianqian Hu, Lily L. Remsing Rix, Bina Desai, Daria Miroshnychenko, Xueli Li, Eric A. Welsh, Bin Fang, Gabriela M. Wright, Neelkamal Chaudhary, Jodi L. Kroeger, Robert C. Doebele, John M. Koomen, Eric B. Haura, Andriy Marusyk, Uwe Rix

## Abstract

Cancer-associated fibroblasts (CAFs) are associated with tumor progression and modulate drug sensitivity of cancer cells. However, the underlying mechanisms are often incompletely understood and crosstalk between tumor cells and CAFs involves soluble secreted as well as adhesion proteins. Interrogating a panel of non-small cell lung cancer (NSCLC) cell lines driven by *EML4-ALK* fusions, we observed substantial CAF-mediated drug resistance to clinical ALK tyrosine kinase inhibitors (TKIs). Array-based cytokine profiling of fibroblast-derived conditioned-media identified HGF-MET signaling as a major contributor to CAF-mediated paracrine resistance that can be overcome by MET TKIs. However, ‘Cell Type specific labeling using Amino acid Precursors’ (CTAP)-based expression and phosphoproteomics in direct coculture also highlighted a critical role for the fibronectin-integrin pathway. Flow cytometry analysis confirmed activation of integrin β1 (ITGB1) in lung cancer cells by CAF coculture. Treatment with pharmacological inhibitors, cancer cell-specific silencing or CRISPR-Cas9-mediated knockout of *ITGB1* overcame adhesion protein-mediated resistance. Concurrent targeting of MET and integrin signaling effectively abrogated CAF-mediated resistance of *EML4-ALK*-driven NSCLC cells to ALK TKIs *in vitro*. Consistently, combination of the ALK TKI alectinib with the MET TKI capmatinib and/or the integrin inhibitor cilengitide was significantly more efficacious than single agent treatment in suppressing tumor growth using an *in vivo EML4-ALK*-dependent allograft mouse model of NSCLC. In summary, these findings emphasize the complexity of resistance-associated crosstalk between CAFs and cancer cells, which can involve multiple concurrent signaling pathways, and illustrate how comprehensive elucidation of paracrine and juxtacrine resistance mechanisms can inform on more effective therapeutic approaches.

## Introduction

Precision therapy with therapeutic antibodies or small molecule drugs can produce high efficacy and relatively low toxicity and has profoundly changed the landscape of cancer therapy.^1^ Such targeted therapies have been particularly impactful in non-small cell lung cancer (NSCLC) leading to a significant improvement in 5-year survival. Anaplastic lymphoma kinase (ALK) tyrosine kinase inhibitors (TKIs), which are approved for NSCLC driven by oncogenic *EML4-ALK* fusions, are amongst the most successful targeted drugs.^2,3^ However, resistance to targeted drugs, including ALK TKIs, remains a major challenge and although multiple mechanisms of resistance have been elucidated, such as secondary point mutations in oncogenes, bypass signaling, histological transformation and epithelial-mesenchymal transition (EMT), in many cases the underlying molecular mechanisms remain elusive.^4,5^ Considering also the heterogeneity of acquired drug resistance mechanisms, there has been increased interest in developing early intervention strategies to target drug-tolerant persister cells,^6^ which derive from drug-sensitive cancer cells that avoid cell death through rapid signaling and subsequent epigenetic adaptations and thus constitute a cell pool from which acquired resistance mechanisms eventually emerge.^7–9^

Cancer cells do not exist in isolation and it is now widely accepted that the tumor microenvironment (TME), in particular cancer-associated fibroblasts (CAFs),^10,11^ which are important components of the TME, can provide critical survival signals causing drug resistance. These signals may furthermore enable cancer cells to withstand the initial impact of targeted drugs and enter a drug-tolerant persister state.^6,12^ Hence, targeting survival crosstalk between CAFs and cancer cells has the potential to enhance the long-term therapeutic outcomes for cancer patients. Notably, recent studies have shown that CAFs not only display phenotypic heterogeneity, but also a high degree of functional complexity regarding signaling mechanisms that can affect cancer cell survival both positively and negatively.^13–15^ Whereas most studies have focused on paracrine signaling by soluble proteins and secreted factors,^16,17^ CAFs also produce various important cell membrane ligands/receptors and extracellular matrix (ECM) proteins. Although some of these proteins are known to strongly modulate the drug responses of cancer cells through adhesion-mediated, i.e. juxtacrine, signaling,^15^ these resistance mechanisms are generally less well understood in the context of TKI resistance compared to paracrine signaling. In order to develop more effective anticancer therapies it is necessary to comprehensively elucidate the complexity of CAF-mediated cancer cell survival mechanisms that encompass both paracrine and juxtacrine signaling. However, concurrent and unbiased interrogation of paracrine and juxtacrine signaling between cancer and stromal cells requires the use of complex technologies to accurately quantify and differentiate signals originating from individual cell types in coculture, which have been successfully applied in only few studies.^18^

In this study, we observed significant CAF-mediated drug resistance of *EML4-ALK* fusion-positive (+) NSCLC cells upon ALK TKI treatment that was elicited by both paracrine and juxtacrine mechanisms. Unbiased and quantitative cell-type specific proteomics and subsequent *in vitro* and *in vivo* functional validation suggested that concurrent targeting of both integrin and HGF-MET signaling is needed to overcome CAF-induced ALK TKI resistance, which can produce novel therapeutic strategies for *EML4-ALK*-positive NSCLC.

## Results

### CAF coculture confers stronger drug resistance to *EML4-ALK*+ NSCLC cells than conditioned medium

To comprehensively determine the impact of CAFs on the drug sensitivity of *EML4-ALK+* NSCLC cells, we assessed 5 established *EML4-ALK+* NSCLC cell lines upon treatment with the Food and Drug Administration (FDA) approved ALK TKIs crizotinib, ceritinib, brigatinib and alectinib. Cell viability was evaluated in cancer cell monoculture, NSCLC cell culture upon exposure to fibroblast conditioned medium (CM), and in coculture with a panel of 4 primary lung CAFs (CAF7, CAF12) and primary (MRC5) or immortalized (Wi38-VA13) normal lung fibroblasts using a live-cell imaging approach based on enumerating the nuclei of NSCLC cells engineered to express red-fluorescent protein in the cell nucleus (**Figure 1A**). The selected fibroblasts are well characterized and have been shown previously to express typical fibroblast marker proteins, such as vimentin (VIM), α-SMA and FAPα, while being devoid of epithelial markers like E-cadherin and cytokeratin.^19^ Employing this system showed that MRC5 lung fibroblasts shielded H3122 and the inherently more resistant CUTO8 and CUTO9 cells against all 4 ALK TKIs (**Figure 1B/C**), particularly at concentrations close to 1 μM, which are readily reached in patient plasma by current first-line ALK TKI.^20–22^ MRC5 CM produced less protective effects implying predominantly cell-contact-dependent mechanisms. Notably, all 4 lung fibroblast cell lines exhibited comparable protective effects irrespective of their origin as lung cancer-associated- or normal fibroblasts (**Figure 1D/E**). Consistent with the findings using MRC5 fibroblasts, coculture with CAF12 conferred significant TKI resistance to 3 of 5 cell lines (**Figure 1D/E/F**, **Figure S1A**). Furthermore, using a trans-well coculture system, which allows for bidirectional paracrine, but not cell-contact-dependent, signaling between fibroblasts and NSCLC cells, showed similar rescue effects compared to CM (**Figure S1B**). This finding suggested that reciprocal signaling via secreted factors alone does not account for the entire extent of drug resistance observed in coculture. In summary, these comparative analyses showed that CAFs conferred significant drug protective effects in several *EML4-ALK+* NSCLC cell lines with varying degrees of protection depending on cancer cell line, fibroblast, TKI and cell culture model. Importantly, coculture elicited stronger resistance than CM alone suggesting resistance mechanisms are mediated by both, soluble and adhesion factors.

**Figure 1.**
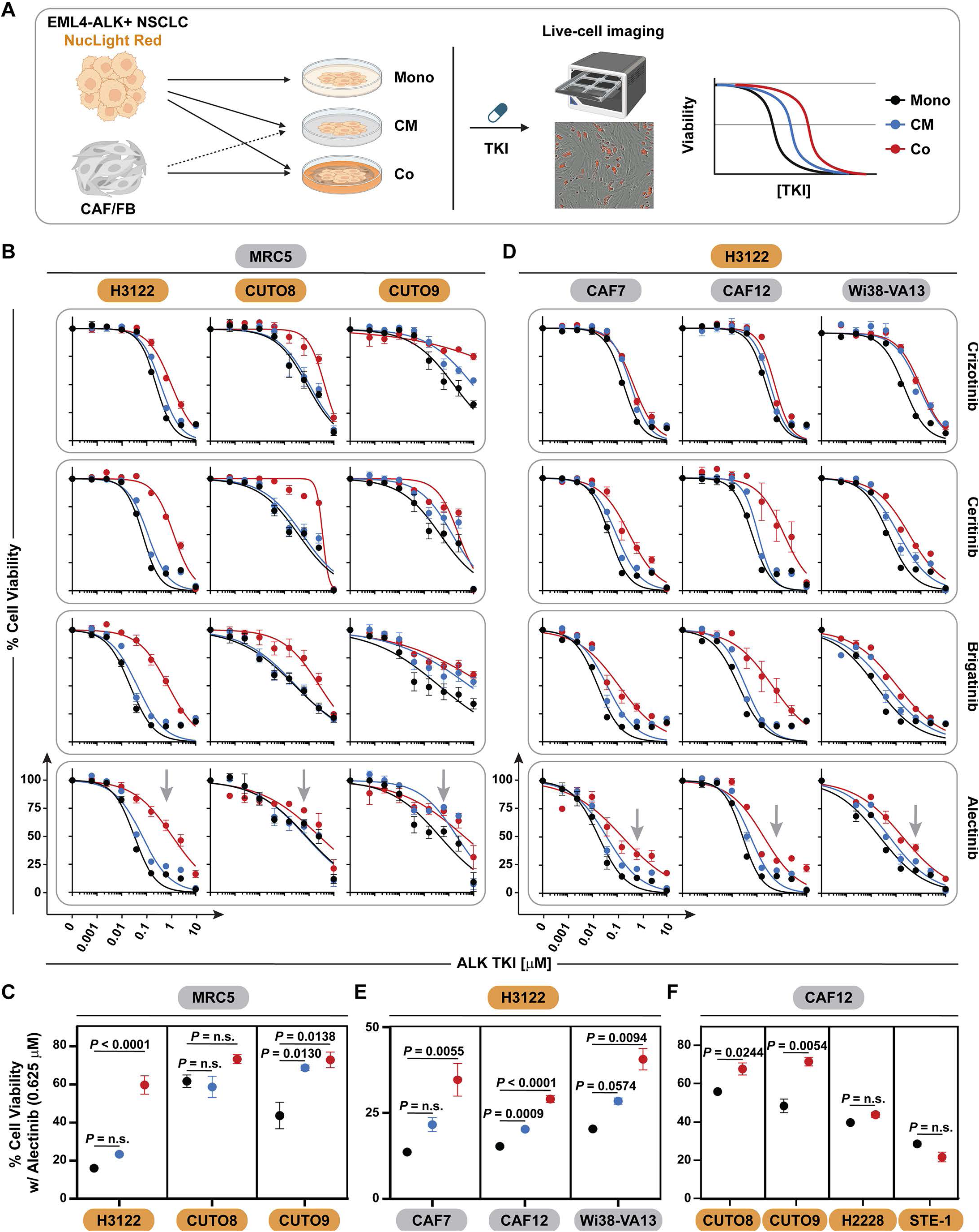
Drug sensitivity modulation of *EML4-ALK*-positive NSCLC cells by fibroblasts. A. Experimental design for drug sensitivity profiling using live-cell imaging of fluorescence-labeled *EML4-ALK*+ NSCLC cells in monoculture (Mono), fibroblast conditioned medium (CM) or physical coculture (Co) with CAFs or normal fibroblasts (FB). **B./C.** Viability of H3122, CUTO8, CUTO9 *EML4-ALK*+ NSCLC cells after 72 hours of treatment with different ALK TKIs in MRC5 CM or coculture. Grey arrows indicate an alectinib concentration of 0.625 μM, which as the closest concentration below the clinically observed Cmax was selected for statistical analysis using one-way ANOVA w/ Holm-Sidak’s multiple comparison correction (**C**). n = 3. **D./E.** Viability of H3122 cells after 72 hours of treatment with different ALK TKIs in CM or coculture of different CAFs/FBs. Grey arrows indicate alectinib concentration selected for statistical analysis (**E**) as described in C. n = 3. **F.** Statistical analysis of viability of different *EML4-ALK*+ NSCLC cells after 72 hours of alectinib treatment in mono- or CAF12 coculture using unpaired t test (see also **Figure S1A**). n = 2.

### HGF-MET signaling is a major driver of CAF-mediated paracrine resistance in *EML4-ALK*+ NSCLC cells

To elucidate the mechanisms underlying the protective role of fibroblast CM on *EML4-ALK-*driven NSCLC cells against ALK TKIs, the secretion profiles of MRC5 fibroblasts, H3122 NSCLC cells and their coculture were analyzed (**Figure 2A**). Analysis of a cytokine array encompassing 80 human proteins, mostly cytokines and growth factors, showed that H3122 secreted predominantly IGFBP-2 and interleukin 8 (IL-8), whereas MRC5 fibroblasts displayed a broader signature of secreted proteins including CCL2, interleukin 6 (IL-6) and IL-8, hepatocyte growth factor (HGF), OPG, TIMP-1 and TIMP-2. Among these, CCL2 and IL-8 are known to promote chemotaxis of immune cells, which would not affect the phenotype in the coculture assay employed here, but may play a role *in vivo*. However, IL-6,^23–25^ OPG,^26,27^ and HGF^17,28–30^ have been previously implicated in cancer cell survival through autocrine or paracrine mechanisms. Considering these prior studies, focus was given to the effects of IL-6, OPG and HGF on *EML4-ALK+* NSCLC cell sensitivity to ALK TKIs. Treatment with recombinant proteins indicated that neither OPG or IL-6 were able to rescue H3122 cells from treatment with various ALK TKIs (**Figure S2A/B**). Notably, whereas addition of recombinant IL-6 to *KRAS*-mutant A549 cells effectively induced downstream STAT3 activation as expected, in H3122 cells IL-6 failed to show such effect (data not shown), indicating a disconnect along the IL-6-STAT3 signaling axis in this cell line. In contrast, HGF, the ligand for the receptor tyrosine kinase MET, has previously been shown to cause resistance of H3122 cells to the preclinical ALK inhibitor TAE684 and lorlatinib.^17,29,31^ As this effect was observed at an HGF concentration of 50 ng/mL, we next determined the concentration of secreted HGF in MRC5 and CAF12 CM, which was found to be between 1.1 and 1.3 ng/mL (**Figure S2C**). Notably, both 1.3 ng/mL as well as 50 ng/mL of recombinant HGF were able to significantly protect H3122 from alectinib treatment (**Figure 2B**), the ALK TKI that is currently most employed in clinical settings. In addition, CAF12 CM-induced alectinib resistance was essentially abrogated by treatment with the selective MET TKI capmatinib suggesting that the majority of the ability of CM to promote drug resistance can be attributed to the action of secreted HGF. Consistently, treatment of H3122 cells with MRC5 CM induced autophosphorylation of MET^Tyr1234/Tyr1235^ and increased phosphorylation of downstream pro-survival proteins, such as extracellular signal– regulated kinase 1/2 (ERK1/2; Thr^202/185^/Tyr^204/187^) and AKT^S473^ (**Figure 2C**), which are critical downstream signals of EML4-ALK^32^ and accordingly were reduced upon alectinib treatment. MRC5 CM partially rescued ERK1/2 and AKT phosphorylation in the presence of alectinib and, conversely, the MET TKI capmatinib abrogated CM-mediated MET and ERK1/2 phosphorylation although this was not equally the case for AKT phosphorylation. Next, we therefore evaluated whether capmatinib can overcome MRC5-mediated resistance to alectinib of H3122 cells with regard to cell viability. Indeed, whereas capmatinib treatment had no effect in monoculture, it completely overcame CM-induced alectinib resistance indicating that HGF is the major functionally relevant component in MRC5 CM pertaining to ALK TKI resistance (**Figure 2D/E**). However, consistent with our previous observations, physical coculture elicited more pronounced alectinib resistance than CM and this resistance was only partially overcome by capmatinib (**Figure 2D/F**). This indicated the presence of mechanisms independent of HGF-MET signaling that significantly contribute to the drug resistance observed in coculture. Taken together, these findings suggest a pivotal role of HGF-MET signaling in CAF-mediated paracrine resistance. However, the partial mitigation of coculture-mediated resistance by MET inhibition also highlights a significant contribution by adhesion factors.

**Figure 2.**
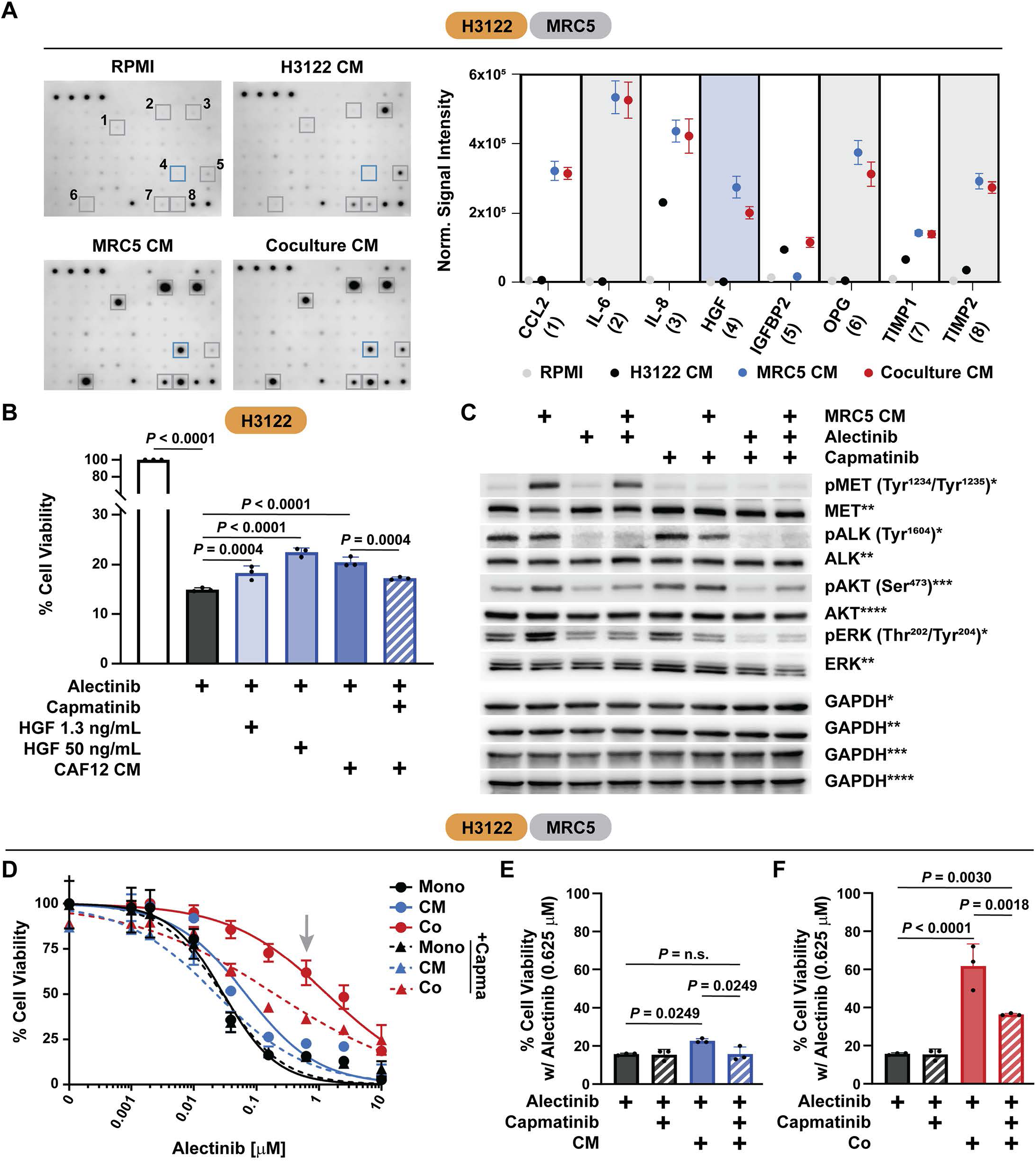
Effects of fibroblast secreted factors on TKI sensitivity of *EML4-ALK*-positive NSCLC cells. A. Cytokine array and quantification of RPMI media, CM from H3122, MRC5, and H3122/MRC5 coculture. n = 2. **B.** Viability of H3122 cells upon treatment with alectinib (1 μM) and capmatinib (0.2 μM) in the presence of HGF or CAF12 CM. *P* values were determined using one-way ANOVA w/ Holm-Sidak’s multiple comparison correction. n = 3. **C.** Immunoblotting for indicated signals in H3122 cells upon treatment with MRC5 CM and/or alectinib (1 μM) and capmatinib (0.2 μM). n = 3. **D.-F.** Viability of H3122 cells upon treatment with alectinib and capmatinib (0.2 μM) in monoculture or in MRC5 CM or coculture. Grey arrow indicates alectinib concentration of 0.625 μM, which was selected for statistical analysis using one-way ANOVA w/ Holm-Sidak’s multiple comparison correction (**E/F**). n = 3.

### Cell-type specific proteomics identifies key role for integrin signaling in CAF coculture- driven resistance

To capture the complexity of cross-communication between cancer cells and CAFs in coculture and discern the respective signaling effects in each cell type, we performed heterocellular phospho- and expression proteomics using a cell-type specific labeling with amino acid precursors (CTAP) approach.^33,34^ This technology utilizes simultaneous stable isotope labeling with d8-D-lysine and diaminopimelate (DAP), respectively. These specific L-lysine precursors are selectively metabolized in only one or the other cell type, which are engineered to express the respective metabolic enzymes, lysine racemase (lyr) and diaminopimelate decarboxylase (DDC). To also determine the signaling changes upon drug treatment, we used isobaric tandem mass tag (TMT) labeling for comparison of alectinib and vehicle (DMSO) treatment. This two-dimensional CTAP- and TMT-labeling-based quantitative proteomics approach enabled analysis of cell-contact dependent signaling both in the presence or absence of ALK TKI treatment (**Figure 3A**). In order to introduce DDC into CAF12 fibroblasts and avoid subsequent senescence during extended metabolic labeling, primary CAF12 cells were immortalized by transduction with human telomerase reverse transcriptase (hTERT) expressing virus. Immortalization of CAF12 did not change expression of fibroblast markers and their ability to cause drug resistance of H3122 cells (**Figure S3A-C**). To further improve cellular growth properties under CTAP labeling conditions, CTAP vectors were enhanced by codon optimization for expression in human cells, incorporation of genes encoding fluorescence proteins, and reassembly into a lentiviral expression systems. Expression of Ddc and Lyr was confirmed by immunoblotting (**Figure S3D/E**). To allow for sufficient cell adaptation upon coculture, CTAP-labeled H3122 and CAF12 cells were cocultured for 24 hours. Alectinib and DMSO treatment were performed for 3 hours to capture immediate signaling responses. Subsequent high resolution liquid chromatography tandem mass spectrometry (LC-MS/MS) analysis detected 4,359 phosphosites and 4,320 lysine-containing peptides with cell-type specific resolution for phospho- and expression proteomics, respectively (**Data S1-4**), which compared favorably with a complex two-dimensional CTAP-labeling study in pancreatic cancer and stellate cells.^18^ Both phospho- and expression proteomics datasets showed little variance across biological replicates (**Figure S3F/G**). As expected, alectinib treatment in monoculture inhibited the phosphorylation of ALK (Tyr^1078^ and Tyr^1096^)^35,36^ and SHP2/PTPN11 (Tyr^542^),^37^ which are important for ALK signaling (**Figure 3B/C**). We also observed upregulation of known compensatory signals, including SRC (Tyr^419^) and EGFR (Tyr^1172^).^38–40^

**Figure 3.**
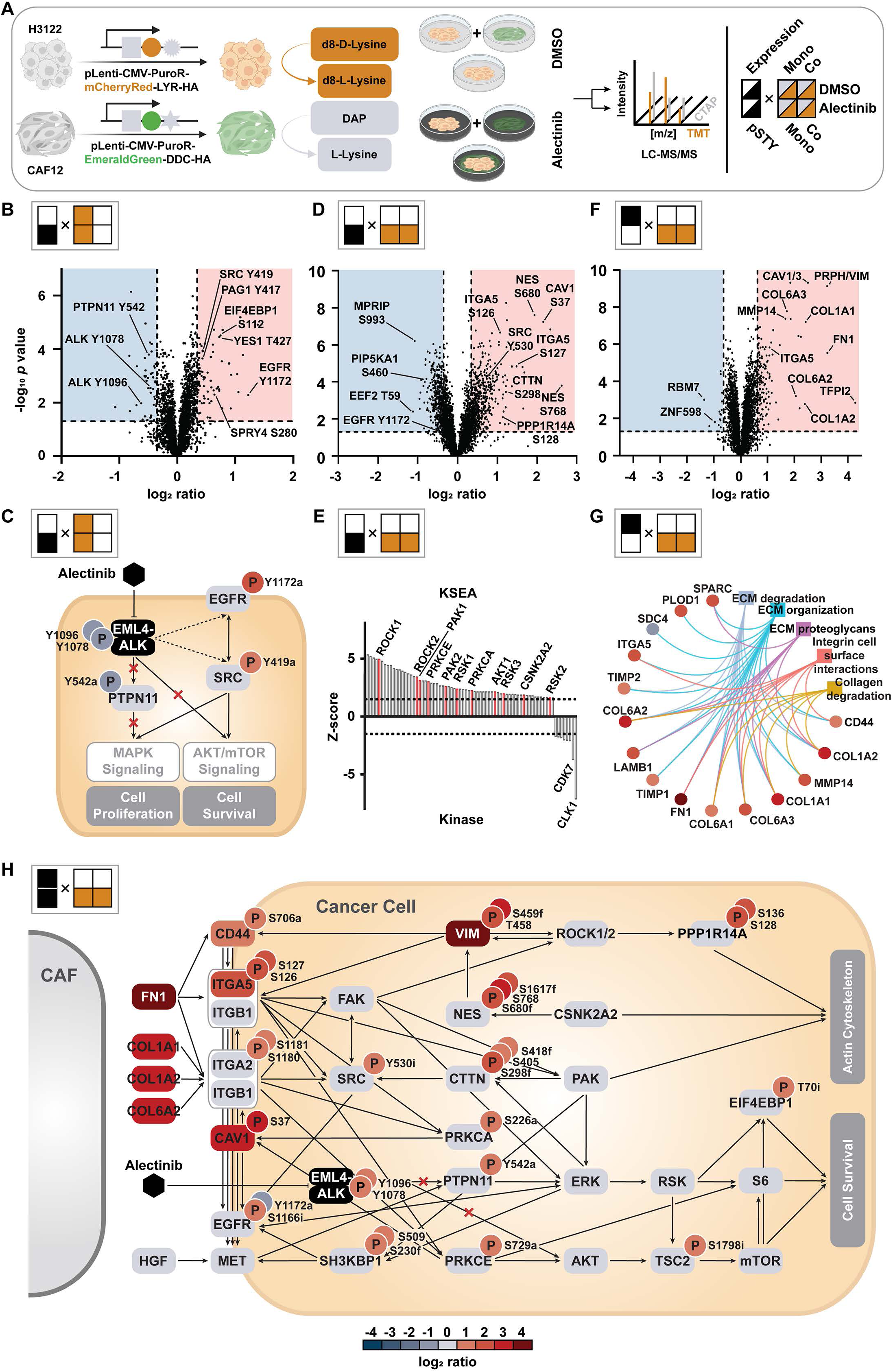
Phospho- and expression proteomics of *EML4-ALK*-positive NSCLC cells in mono- and coculture with CAFs. A. Experimental design for quantitative global phospho- and expression proteomics of H3122 and hTERT-CAF12 cells in mono- and coculture upon treatment with alectinib (1 μM) or DMSO using a CTAP- and TMT-based dual labeling strategy. **B.** Global phosphorylation changes in H3122 cells upon treatment with alectinib compared to DMSO in monoculture. Significantly regulated signals are indicated by highlighted areas (up: red; down: blue). **C.** Key pathway effects in H3122 cells based on phosphoproteomics analysis from B. **D.** Global phosphorylation changes in H3122 cells upon treatment with alectinib in coculture compared to monoculture. **E.** KSEA-predicted kinase activity changes in H3122 cells based on phosphoproteomics analysis from D. **F.** Protein expression changes in H3122 cells upon treatment with alectinib in coculture compared to monoculture. **G.** Pathway enrichment by overexpressed proteins from expression analysis from F. **H.** Cartoon of major signaling changes in H3122 cells upon CAF12 coculture and alectinib treatment from integration of phospho- and expression proteomics data.

These results indicated that CTAP-based proteomics can capture functionally relevant proteome alterations and signaling changes. Besides the drug response in a cell-autonomous manner, the cancer cell proteome was further modulated by cell interaction with CAFs (**Figure 3D-G**). Intriguingly, the changes induced by coculture were yet more pronounced than those induced by alectinib treatment (**Figure S4A**). Relative to monoculture, coculture with CAFs dampened compensatory survival signals by suppressing activating phosphorylation of EGFR (Tyr^1172^) and inducing inhibitory phosphorylation of SRC (Tyr^530^). Interaction with CAFs also substantially modulated phosphorylation of multiple adhesion signaling-related proteins, such as CTTN (Ser^298^), NES (Ser^768^), ITGA5 (Ser^126/7^) and CAV1 (Ser^37^) (**Figure 3D**). Kinase-Substrate Enrichment Analysis (KSEA)^41^ suggested activation of the adhesion-relevant kinases ROCK1/2, PAK1/2, PRKCA/E as well as pro-survival kinases RSK1/2/3, AKT1 and CSNK2A2 (**Figure 3E**)., Whereas most of these signals represented changes in phosphorylation, expression proteomics analysis indicated that some may have been primarily upregulated by increased protein abundance, such as VIM, CAV1 and ITGA5 (**Figure 3F**). It also highlighted elevated expression of integrin-associated ECM ligands fibronectin (FN1) and various collagens. Notably, abundance more so than phosphorylation is relevant for downstream signaling by these proteins. Consistently, pathway analysis of proteins that were differentially expressed in alectinib-treated coculture suggested activated ECM-integrin-related signaling (**Figure 3G**). Furthermore, many of the observed changes, such as dampened EGFR signaling and activated integrin signaling, were consistent with coculture-induced proteome changes in CAF12 fibroblasts (**Figure S4B-E**), particularly reduced and increased expression of the secreted proteins AREG and FN1, respectively (**Figure S4E**). Incorporating protein expression and phosphorylation changes induced in alectinib-treated cancer cells by interaction with CAFs generated an integrated signaling network map (**Figure 3H**). This map illustrated the interconnectivity of signaling alterations and that CAFs compensated for much of the inhibitory effect of alectinib on EML4-ALK and the downstream pro-survival ERK/RSK and AKT/mTOR signaling pathways by activating ECM-integrin signaling. Specifically, multiple ECM ligands were enriched in coculture leading to overactivation of integrins, which function as heterodimers of beta and alpha subunits. Thus, the overexpression of ITGA5 and the hyperphosphorylation of ITGA2 and ITGA5 indicated converged overactivation of ITGB1 heterodimers, the only binding partner for these two alpha subunits. In addition, FN1 can also activate CD44,^42,43^ which is further facilitated by intracellular VIM and can stimulate integrin-mediated adhesion.^44,45^ VIM furthermore activates ROCK1/2, PPP1R14A and ITGA5/ITGB1 heterodimers. Activated integrin receptors can also interact with other receptors, including EGFR^46^ and MET.^47^ Intriguingly, we observed increased activation of PTPN11, which was not due to EML4-ALK considering inhibition of the latter by alectinib. PTPN11 phosphorylation may be the result of MET activation directly via integrin heterodimers and/or via HGF,^48^ which would be consistent with secretion of HGF by these CAFs (see **Figure S2C**). In summary, quantitative cell-type specific phospho- and expression proteomics highlighted upregulation of multiple cell survival-associated ECM-integrin-related adhesion signals upon coculture of *EML4-ALK*+ NSCLC cells with CAFs. Importantly, these signals converged on ITGB1, which emerged as a potential target for neutralizing coculture-mediated protective effects to ALK TKIs.

### ECM-integrin signaling causes coculture-mediated resistance by CAFs

Given the robust upregulation of ECM-integrin signaling in coculture, we hypothesized that inhibiting this pathway suppresses coculture-mediated rescue in response to drug treatment. ITGB1 is the most prevalent subunit in integrin heterodimers and can interact with 12 of the 18 alpha subunits.^49^ We therefore first quantified the activation level of ITGB1 and the key EML4- ALK downstream survival signal ERK (Thr^202^/Tyr^204^) in H3122 monoculture and in coculture with CAF12 using flow cytometry, both in the absence and presence of alectinib (**Figure 4A**). Consistent with the observations from CTAP-based proteomics, coculture strongly activated ITGB1 irrespective of alectinib treatment. Importantly, the pan-integrin antagonist cilengitide, a peptidomimetic that competes for the arginine-glycine-aspartic acid (RGD) sequence critical for integrin-ligand binding,^50^ reduced this effect to levels seen in monoculture. Phosphorylation of ERK was, as expected, downregulated by alectinib. This effect was less pronounced in coculture. However, addition of cilengitide reduced ERK phosphorylation in coculture more markedly. Consistently, silencing of *ITGB1* in H3122 cells by RNA interference partially overcame CAF12 coculture-mediated resistance to alectinib (**Figure 4B**). As FN1 is a major activator of integrin signaling and was one of the most strongly upregulated proteins by CAFs in coculture according to CTAP-based proteomics, we investigated the impact of FN1 as a surrogate for CAF coculture on H3122 cell viability and intracellular signaling upon alectinib treatment. FN1 significantly protected H3122 cells in the presence of alectinib, yet this effect was abrogated by cilengitide (**Figure 4C**). Correspondingly, FN1 partially rescued the downstream pro-survival signals ERK (Thr^202^/Tyr^204^), AKT (Ser^473^), and S6 (Ser^235^/Ser^236^). This was consistent with increased autophosphorylation of FAK (Tyr^397^), which acts downstream of integrins, but upstream of MAPK and AKT/mTOR signaling. Notably, co-treatment with cilengitide overcame FN1-mediated rescue leading to almost complete abrogation of these signals (**Figure 4D**). Collectively, these results suggested that CAF coculture activated ITGB1 and downstream pro-survival signals via FN1, which resulted in partial protection from alectinib treatment that can be overcome by integrin targeting.

**Figure 4.**
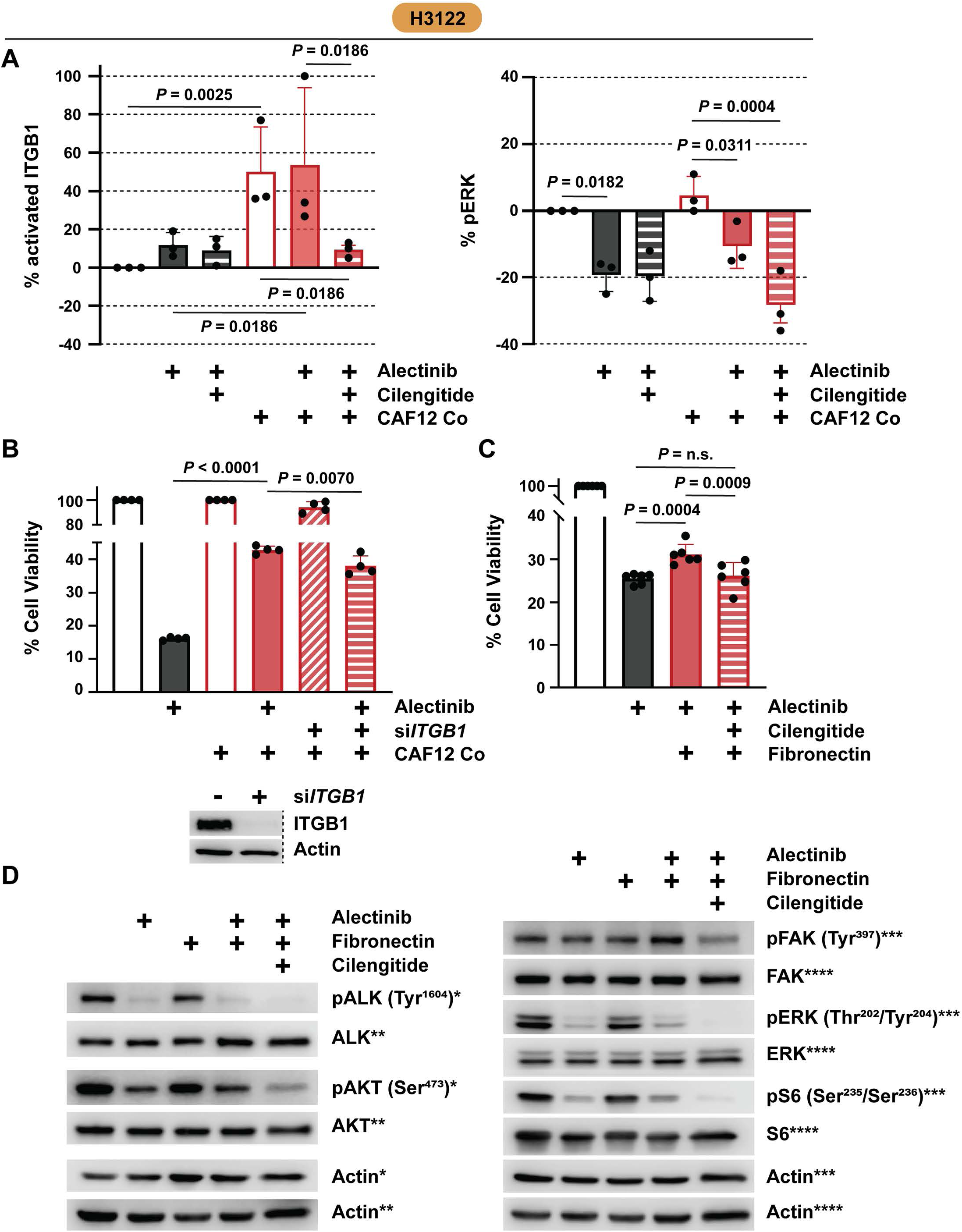
Effects of targeting integrin signaling on coculture-mediated drug resistance. A. ITGB1 and ERK activation levels in H3122 cells in mono- or coculture with CAF12 upon alectinib treatment (1 μM, 3 hours) as determined by flow cytometry. *P* values were determined using one- way ANOVA w/ Holm-Sidak’s multiple comparison correction. n = 3. **B.** Viability of H3122 cells in mono- or coculture with CAF12 upon treatment with si*ITGB1* (pool) or non-targeting siNT control for 96 hours, and alectinib (1 μM) for 72 hours. *P* values were determined using one-way ANOVA w/ Holm-Sidak’s multiple comparison correction. n = 4. **C.** Viability of H3122 cells treated for 72 hours with alectinib (1 μM) and cilengitide (1 μM) with or without fibronectin coating (2.5 μg/cm^2^). *P* values were determined using one-way ANOVA w/ Holm-Sidak’s multiple comparison correction. n = 6. **D.** Immunoblotting for indicated signals in H3122 cells upon coating with fibronectin (2.5 μg/cm^2^) and/or treatment with alectinib (1 μM) and cilengitide (1 μM). Asterisks denote signals from same membranes. n = 3.

### Combinatorial inhibition of ITGB1 and MET overcomes CAF-induced rescue effect

Cytokine analysis and CTAP-proteomics identified that MET and ITGB1 signaling concurrently mediate the CAF-driven rescue effects. Consistently, CRISPR-Cas9 knockout of *ITGB1* or MET inhibition by capmatinib each significantly reduced, but only partially overcame CAF12-mediated alectinib resistance of H3122 cells (**Figure 5A/B**). However, combined *ITGB1* knockout and MET inhibition completely abrogated coculture-induced resistance at physiologically relevant concentrations and decreased cell viability to monoculture levels. *ITGB1* knockout also significantly reduced coculture-induced alectinib resistance in CUTO9 cells (**Figure S5A**). In strong agreement with the CRISPR-based knockout results, dual pharmacological targeting of ITGB1 and MET by cilengitide and capmatinib, respectively, completely abolished CAF12 coculture-driven alectinib resistance of H3122 cells, whereas single drug treatment only resulted in partial re-sensitization (**Figure 5C/D**). Similarly, combination of capmatinib with a second integrin antagonist, GLPG0187, corroborated these observations (**Figure S5B/C**). Furthermore, combined treatment of cilengitide and capmatinib also completely overcame CAF12 coculture-mediated alectinib resistance in the inherently less ALK TKI-sensitive CUTO8 and CUTO9 cells (**Figure S5D/E**). Notably, although CUTO8 cells showed less sensitivity to capmatinib in mono- or coculture, the addition of capmatinib to cilengitide yielded still more pronounced effects than cilengitide treatment alone in mitigating alectinib resistance. Taken together, these results suggest that CAF-mediated protection of *EML4-ALK*+ NSCLC cells from ALK TKIs in coculture can be overcome by and requires combined targeting of ITGB1 and MET.

**Figure 5.**
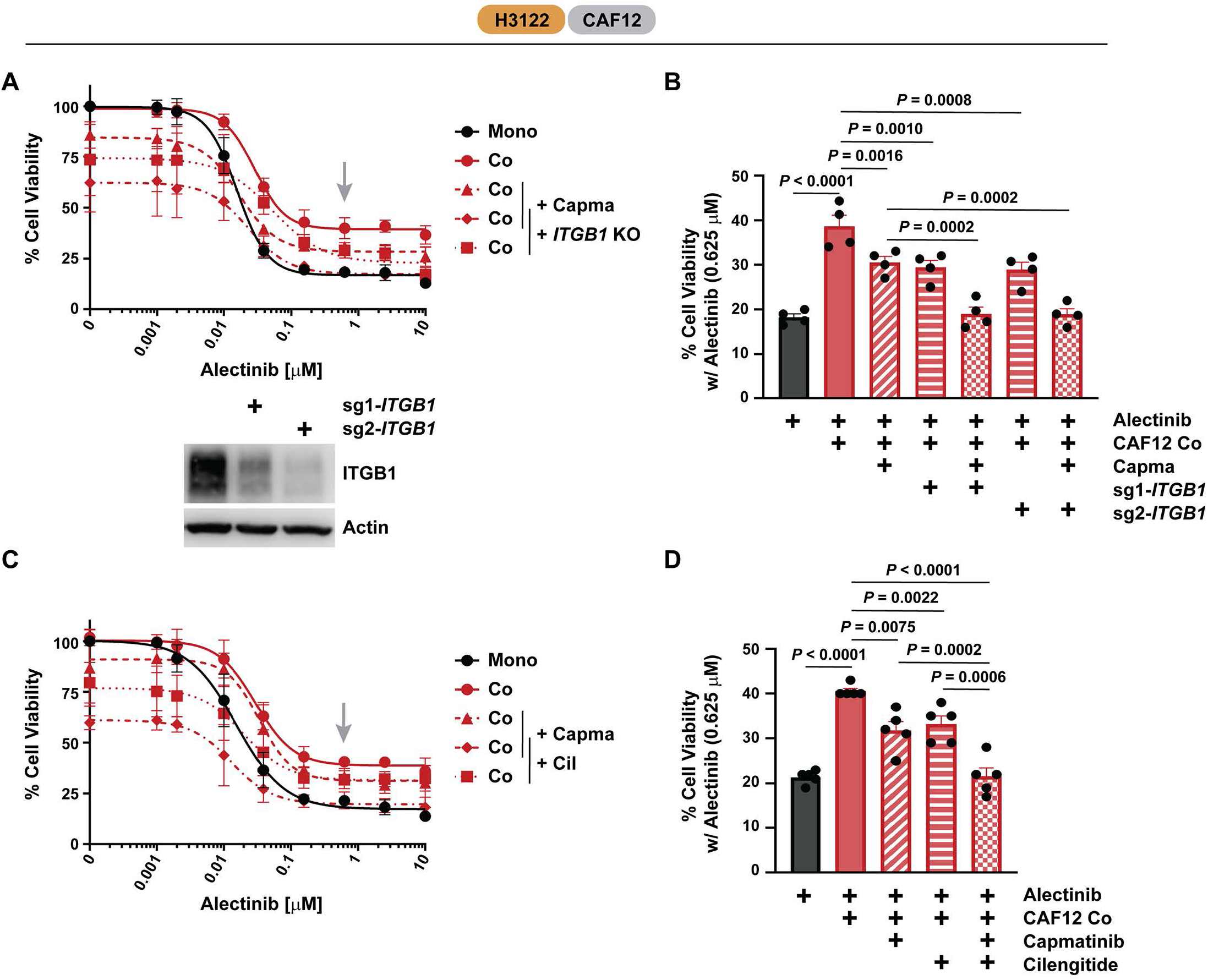
Combined targeting of coculture-induced ITGB1- and MET-mediated resistance in H3122 cells. A./B. Viability of H3122 cells upon CRISPR-mediated knockout of *ITGB1* and treatment with alectinib and capmatinib (0.2 μM) in mono- or CAF12 coculture. Grey arrow indicates alectinib concentration of 0.625 μM, which was selected for statistical analysis using one-way ANOVA w/ Holm-Sidak’s multiple comparison correction (**B**). n = 4. **C./D.** Viability of H3122 cells upon treatment with alectinib, cilengitide (1 μM) and capmatinib (0.2 μM) in mono- or CAF12 coculture. Grey arrow indicates alectinib concentration of 0.625 μM, which was selected for statistical analysis using one-way ANOVA w/ Holm-Sidak’s multiple comparison correction (**B**). n = 5.

### Targeting ITGB1 and MET improves EML4-ALK+ tumor response

Considering the necessity to target both ITGB1 and MET in order to fully abrogate CAF-mediated rescue effects in *EML4-ALK*+ NSCLC cells, we next evaluated the *in vivo* therapeutic efficacy of combinatorial treatment compared to alectinib treatment alone. To this end, tumor growth was monitored in an *EML4-ALK*+ allograft mouse model,^51^ which features an intact tumor microenvironment, following different single and combination drug regimens over a 4-week period. Both the dual combination regimens of alectinib with capmatinib or cilengitide, and the triple regimen, were well tolerated, as indicated by the stable mouse body weights throughout the treatment duration (**Figure S5F**). As expected, single agent capmatinib or cilengitide treatment did not show any effect, whereas alectinib inhibited tumor growth. Notably, the combination regimens exhibited superior anti-tumor effects compared to alectinib alone leading to significantly enhanced tumor shrinkage (**Figure 6A-C**). Intriguingly, immunohistochemistry (IHC) analysis revealed that the activating and proliferation-inducing phosphorylation of S6 (Ser^235^/Ser^236^) was increased upon exposure to alectinib alone, potentially as a compensatory signal upon ALK inhibition. However, this upregulation of S6 phosphorylation was suppressed by cilengitide combination therapy, most substantially by the combined addition of cilengitide and capmatinib (**Figure 6D**). This effect was consistent with our previous observation *in vitro* that the combined treatment of alectinib and clilengitide strongly reduced S6 phosphorylation in the presence of FN1 (see **Figure 4D**).

**Figure 6.**
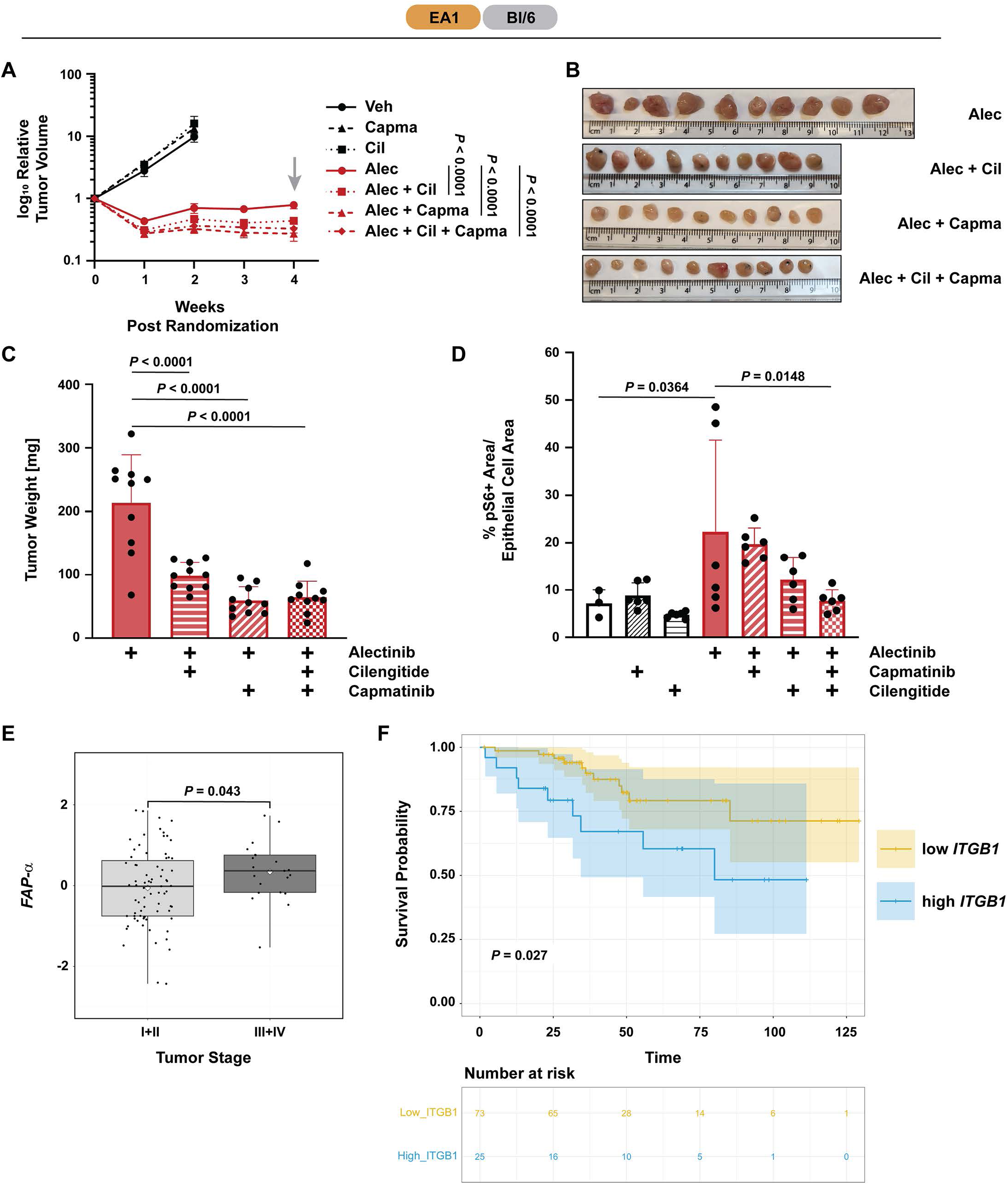
Impact of expression and targeting of ITGB1 in NSCLC tumors. A.. Relative tumor volumes of mEA1 mouse allografts in Bl/6 treated with vehicle control (Veh; 0.5% hydroxypropyl methylcellulose/0.1% Tween 80), alectinib (Alec; 20 mg/kg; p.o.), capmatinib (Capma; 3 mg/kg; p.o.), cilengitide (Cil; 5 mg/kg; i.p.) or various combinations of these. Grey arrow indicates 4 week experiment endpoint, which was selected for statistical analysis. *P* values were determined using two-way ANOVA w/ Holm-Sidak’s multiple comparison correction. n = 10 (exception: Veh/Capma/Cil; n = 6). **B.** Images and sizes of harvested tumors from panel A. **C.** Weights of harvested tumors from panel A. *P* values were determined using one-way ANOVA w/ Holm-Sidak’s multiple comparison correction. **D.** Immunohistochemistry-based quantification of pS6- positive epithelial cell area relative to the total epithelial cell area across different treatment arms at endpoint. n = 6. *P* values were determined using one-way ANOVA w/ Holm-Sidak’s multiple comparison correction. **E./F.** Association analysis of *FAP-α* expression levels and tumor stage (**E**) and survival analysis stratified by low and high expression levels of *ITGB1* (**F**) using *EGFR*-mutant, *ROS1-* or *ALK-*fusion-positive NSCLC patient data from the ‘oncosg’ dataset. *P* values were determined by t test.

To further assess the significance of CAFs in the progression of *EML4-ALK*+ NSCLC, we aimed to investigate the association between CAF composition and tumor stages of these patients using publicly available datasets. Due to the limited sample size of *EML4-ALK*+ NSCLC patients, NSCLC cases driven by *ALK* or *ROS1* fusions or *EGFR* mutations were pooled together for further analysis. As a surrogate for CAFs, the mRNA expression levels of fibroblast activation protein α (FAP-α), a widely used marker for fibroblasts, were used as an indicator for CAF composition in a patient sample. This analysis revealed that higher FAP-α expression levels were associated with more advanced cancer stages in these patients (**Figure 6E**). Furthermore, within the same dataset of *EGFR/ROS1/ALK*-positive NSCLC patient samples, higher mRNA expression of *ITGB1* was associated with significantly worse survival probability (**Figure 6F**). In summary, these results highlighted that targeting integrin and MET signaling in combination with ALK TKIs can overcome CAF-mediated survival signaling for *EML4-ALK*+ NSCLC cells *in vivo* and that this strategy may have significant potential also in NSCLC patient tumors to improve clinical outcomes.

## Discussion

Cancer-associated fibroblasts (CAFs) can have a major impact on the drug sensitivity of cancer cells.^15^ However, due to the diverse challenges of working with CAFs, examples of studies that comprehensively interrogate the complex and dynamic signaling crosstalk between cancer cells and CAFs in an unbiased way, such as a pioneering study in pancreatic cancer and stellate cells,^18^ are rare. Here, we describe use of an adaptation of cell-type specific labeling with amino acid precursors (CTAP) to perform quantitative phospho- and expression proteomics on *EML4-ALK*+ NSCLC cancer cells in physical coculture with lung CAFs, which allowed for concurrent evaluation of both paracrine and juxtacrine signaling mechanisms. This approach elucidated the combined role of survival signaling by integrin β1 (ITGB1) heterodimers and MET in mediating resistance to clinical ALK TKIs. This is consistent with previous studies that illustrated the role of HGF-MET signaling for resistance to ALK TKIs in NSCLC and to other drugs in different cancers.^17,29,52^ However, a concurrent and equally pronounced role for integrin signaling in eliciting CAF-mediated ALK TKI resistance has not been described. Notably, *ITGB1* expression correlated with poor patient survival in the clinics. Whereas these effects were pronounced and observed across different *EML4-ALK*+ NSCLC cells and CAFs, given the broad heterogeneity of CAFs and the complexity of cancer cell/CAF crosstalk,^31,53–55^ it is possible that additional signaling proteins and pathways further modulate ALK inhibitor responses in different cancer cell line/CAF pairs. This is likely to be relevant for other drugs in different tumor types, as well, and may even reverse drug resistance as recently shown in some cases.^19,56,57^ However, it is notable that HGF-MET and ITGB1 signaling have been separately identified to cause BRAF inhibitor resistance in melanoma,^17,28,58^ and may therefore work cooperatively here, too. Importantly, CAF-induced ITGB1-FAK signaling was found to promote *BRAF*-mutant melanoma persister cell survival upon BRAF inhibitor treatment.^58^ Consistently, although they focused on tumor-intrinsic mechanisms and did not investigate the contribution from integrins or CAFs, the importance of FAK-YAP1 signaling for the survival of persister cells was recently demonstrated also in NSCLC,^59^ which is in excellent agreement with our observations here as FAK acts downstream of integrins. However, CAFs caused resistance also to lorlatinib^31^, ceritinib and brigatinib, which besides being ALK TKIs also potently inhibit FAK (and PYK2).^60–64^ This seeming discrepancy may be explained by the fact that integrins do not exclusively signal through FAK (or PYK2), but can also utilize other downstream pathways, e.g. via integrin-linked kinase (ILK), SRC, talin and other molecules.^49,65^ As discussed furthermore earlier, CAFs promote *EML4-ALK*+ NSCLC cell survival in part through HGF-MET signaling, thereby circumventing FAK inhibition. Moreover, there is marked signaling crosstalk between integrins and MET as ITGB1 can activate MET in an HGF-independent fashion via CD44 and fibronectin-activated α5β1 integrin heterodimers.^44,47,66,67^ Adding complexity to the matter, neither is MET only activated by ITGB1 (but also by HGF), nor is ITGB1 restricted to signal only through MET (but also through FAK, ILK, etc.). This crosstalk illustrates that in order to fully abrogate CAF-mediated rescue of *EML4-ALK*+ NSCLC cells from ALK TKIs, it is necessary to target both pathways simultaneously, which is consistent with our observations that MET and ITGB1 inhibitors cannot fully replace one another, but act in a cooperative fashion.

The observed signaling complexity poses significant challenges for translating these findings as triplet drug combinations are difficult to implement in the clinics due to logistic/business-related hurdles and a heightened risk of toxicity. Our data showed good tolerability of the alectinib/capmatinib/cilengitide combination in mice, but another possible solution to address this challenge may be to harness the serendipitous polypharmacology of some clinically approved TKIs.^68^ For instance, combining the dual ALK/FAK TKIs lorlatinib, ceritinib or brigatinib with selective MET TKIs, such as capmatinib or tepotinib, or combining the dual ALK/MET TKI crizotinib with an integrin inhibitor may elicit efficacy similar to a triple drug combination for these targets. Notably, a study aiming to enhance ALK TKI efficacy in *EML4- ALK*+ NSCLC by targeting tumor cell-intrinsic integrin β3 overexpression demonstrated that the combination of crizotinib and cilengitide was tolerated in a mouse model.^69^ To further mitigate potential toxicity concerns, it may be even feasible to implement sequential switching regimens of these combinations, which may allow to benefit from the more potent inhibition of ALK by the next-generation ALK TKIs lorlatinib and brigatinib while maintaining intermittent MET inhibition through crizotinib. Thus, there may be tangible avenues for clinical translation of the findings presented here although further studies are required to select the optimal therapeutic strategy and carefully evaluate its feasibility in clinical trials.

In summary, in this study we used an innovative quantitative cell-type specific functional proteomics approach that enabled dissecting the complex signaling mechanisms, by which cancer-associated fibroblasts (CAFs) confer drug resistance in *EML4-ALK*+ NSCLC cells, and identified both juxtacrine and paracrine signaling through integrin β1 (ITGB1) and MET to cooperatively rescue cancer cells from ALK TKIs. The superior efficacy observed with the combinatorial inhibition of ITGB1 and MET, in addition to standard of care drugs, underscored the importance of concurrent integrin and MET signaling in modulating the therapeutic response to ALK TKIs in EML4-ALK+ NSCLC and illustrated that in order to overcome CAF-mediated resistance, it is necessary to target multiple survival signaling pathways simultaneously. However, considering the inherent polypharmacology of some clinical ALK TKIs, these findings can rationally inform translational combination studies with increased efficacy and potentially manageable safety profiles.

## Materials and Methods

### Cell culture and reagents

H3122, H2228, STE1 and WI38-VA13 cell lines were obtained from the Moffitt Cancer Center Lung Cancer Center of Excellence Cell Line Core. CUTO8 and CUTO9 cell lines were generated by Dr. Robert Doebele at the University of Colorado as described previously.^70^ MRC5 lung fibroblasts were purchased from the American Type Culture Collection (ATCC). CAF7 and CAF12 were generated as previously described.^71^ All cell lines were tested negative for mycoplasma and authenticated via short tandem repeat analysis. All cells were cultured in RPMI 1640 medium supplemented with 10% fetal bovine serum at 37°C in a humidified atmosphere containing 5% CO2. Nuclear red labeling of cancer cells was achieved using Incucyte® Nuclight Red Lentivirus reagent (Sartorius #4476) according to the manufacturer’s instructions with modifications described previously.^19^

pENTR1A no ccDB (w48-1) and pLenti CMV Puro DEST (w118-1) were gifts from Eric Campeau & Paul Kaufman (Addgene plasmid #17398, http://n2t.net/addgene:17398, RRID:Addgene_17398; Addgene plasmid #17452, http://n2t.net/addgene:17452, RRID:Addgene_17452).^72^ Lenti_EFS-Cas9-P2A-HygR was a gift from Hyongbum Kim (Addgene plasmid #164134, http://n2t.net/addgene:164134; RRID:Addgene_164134).^73^ LRG2.1 was a gift from Christopher Vakoc (Addgene plasmid #108098, http://n2t.net/addgene:108098, RRID:Addgene_108098).^74^ pLV-hTERT-IRES-hygro was a gift from Tobias Meyer (Addgene plasmid #85140; http://n2t.net/addgene:85140; RRID:Addgene_85140)^75^ and packaged with a 3^rd^ generation lentivirus packaging system (ABM, LV053). CAF12 cells were transduced with hTERT plasmid-containing lentivirus followed by selection with 50 μg/mL hygromycin B (EnzoLifeSciences, #ALX-380-306-G001). GLPG0187 was purchased from Cayman Chemical (#21792).

### Cell viability assay upon CM, trans-well and physical coculture

Conditioned media (CM) were generated as described before.^19^ For viability determination in CM, cells were plated in a 1:1 mixture of normal growth medium (RPMI + 10% FBS) and CM. Trans- well cultures were set up using permeable support (Corning® HTS Transwell® 96 well permeable supports, #3381), where 1,000 cancer cells were seeded in the bottom wells and 1,000 CAF cells were seeded on permeable trans-wells the day before drug addition. Physical coculture was set up by seeding 1,000 cancer cells plus 1,000 CAFs into the same well of a 384-well plate (Corning, #3764) the day before drug treatment to allow 24 hours of coculture. For monoculture controls, the same number of labeled *EML4-ALK* fusion NSCLC cells were plated as in coculture (1,000 cells). Following 72 hours of drug treatment, cell viability was ascertained by quantifying fluorescently labeled red nuclei (red object count per well) on the IncuCyte live-cell analysis system (Essen BioScience, instrument/software versions S3 or SX5). Data were subsequently analyzed utilizing GraphPad Prism.

### Cell viability rescue assay upon HGF, IL-6 and OPG treatment

Human recombinant HGF (PeproTech Inc, #100-39H), IL-6 (Peprotech, #200-06), and OPG (RnD system, # 6945) were reconstituted according to the manufacturer’s instructions. HGF was added at concentrations of 1.3 ng/mL and 50 ng/mL at the time of cell seeding. IL-6 and OPG were added at 20 ng/mL and 100 ng/mL, respectively. For monoculture controls, cells were seeded in regular media (RPMI + 10% FBS). The cell viability was determined after 72 hours of drug treatment as described before.

### Cytokine array

Regular media vs CM from H3122, MRC5 and coculture of both were collected after 24 hours of monoculture or coculture. The cytokine array assay was performed in 2 biological replicates and the results were quantified according to the manufacturer’s instructions (RayBiotech, #AAH-CYT- 5-8). Images were acquired using an Odyssey FC Imager (LI-COR) and quantified using Image Studio Lite Ver 5.2.

### HGF ELISA

CM was collected after 24 hour of cell culture. The concentrations of HGF were then determined using the human HGF ELISA kit (Sigma-Aldrich, #RAB0212) according to the manufacturer’s instructions.

### Immunoblotting

Cell lysates were separated by SDS-PAGE and immunoblotting was performed with primary antibodies against β-actin (Sigma, A5441), total ALK (Cell Signaling, #3633), pALK (Tyr^1604^) (Cell Signaling, #3341), total MET (Cell Signaling, #8198), pMET (Tyr^1234/1235^) (Cell Signaling, #3077), total AKT (Cell Signaling, #9272), pAKT (Ser^473^) (Cell Signaling, #4060), total ERK (Sigma, #M5670), pERK (Thr^202^/Tyr^204^) (Cell Signaling, #4370), GAPDH (Proteintech, # 60004), HA-tag (Cell Signaling, #3724), ITGB1 (Cell Signaling, #9699), hTERT (Rockland, #600-401-252), αSMA (Abcam, #ab32575), E-cadherin (Cell Signaling, #3195), S6 (Cell Signaling, #2217), pS6 (Ser^235^/Ser^236^) (Cell Signaling, #4858), FAK (Cell Signaling, #13009) and pFAK (Tyr^397^) (Cell Signaling, #8556). Secondary antibodies were anti-rabbit and anti-mouse (GE Healthcare). Images were acquired using an Odyssey FC Imager (LI-COR) and quantified using Image Studio Lite Ver 5.2. For the immunoblotting upon drug and CM treatment, cells were seeded at 0.6*10^6^ cells/well in 6-well plate on day 0. On day 1, cells were harvested after 1 hour of drug +/- CM treatment and processed by SDS-PAGE and immunoblotting.

### Transgenic expression of L-lysine biosynthesis gene

C-terminal HA- and Kdel-tagged *Lyr* and *Ddc* expression vectors were kindly provided by Gauthier et al.,^33^ and codon-optimized for expression in human cells. Next, a T2A sequence was cloned into vectors upstream of *Lyr* and *Ddc*. The optimized and HA-tagged T2A-*Lyr* and T2A-*Ddc* were cloned into the EcoRI/EcoRV sites of the pENTR1A vector (Addgene, #17398). G-blocks for human codon-optimized *mCherry* and *Emerald green* were synthesized (IDT) and cloned via BamHI/EcoRI sites into pENTR1A-T2A-LYR/DDC vectors, respectively. Lastly, these constructs were recombined using LR-Clonase (Gateway) into the pLenti-CMV-Puro-DEST vector (Addgene, #17452) resulting in the final expression vectors of pLenti-CMV-Puro-mCherry-T2A- Lyr-HA-Kdel and pLenti-CMV-Puro-Emerald-T2A-Ddc-HA-Kdel as validated by sequencing. Lentivirus particles carrying these expression plasmids were generated using a 3^rd^ generation virus packaging kit (ABM LV053) by transfecting HEK293T cells. Virus was concentrated and titered. H3122 or immortalized CAF12 cells were infected by the lentivirus containing the *Lyr* or *Ddc* expression vectors as described above. Infected cells were selected with 2 μg/mL puromycin for 7 days. Fluorescence of the infected H3122 or CAF12 was monitored by microscopy and validated by flow cytometry. Expression of Lyr and Ddc were detected by immunoblotting for HA.

### CTAP labeling for proteomics

Lyr-H3122 cells were continuously grown in SILAC RPMI (Fisher Scientific, #89984) in the presence of 4 mM ‘heavy’ D-lysine (D-lysine-3,3,4,4,5,5,6,6-d8 2 HCl) (>98% purity, C/D/N Isotopes Inc., D6367). Immortalized Ddc-CAF12 were continuously grown in SILAC media supplemented with 10 mM unlabeled (‘light’) DAP. SILAC RPMI media were additionally supplemented with 10 pM insulin (Sigma, I0516), 1.15 mM L-arginine (Sigma, A8094), and 10% dialyzed FBS (Fisher Scientific, #88440). The cells were cultured separately to reach the heavy label incorporation rate of 97% as validated by MS. hTERT-CAF12 and H3122 cells were then mono- or cocultured for 24 hours followed by DMSO or 1 μM alectinib treatment for 3 hours before cell collection in PBS supplemented with sodium vanadate.

### Proteomics sample preparation and LC-MS/MS analysis

Cell pellets were lysed in denaturing buffer containing 8 M urea, 20 mM HEPES (pH 8.0), 1 mM sodium orthovanadate, 2.5 mM sodium pyrophosphate and 1 mM β-glycerophosphate. the Protein concentrations were determined by Bradford assay. Aliquots of 200 μg were prepared for global phosphorylation/expression analysis. Disulfide bonds were reduced with 4.5 mM DTT at 60°C for 30 minutes and peptides were alkylated with 10 mM iodoacetamide for 20 minutes in the dark at room temperature. Lys-C digestion was carried out at room temperature overnight with enzyme-to-substrate ratio of 1:20. Digested peptides were acidified with aqueous 1% trifluoroacetic acid (TFA) and desalted with C18 Sep-Pak cartridges according to the manufacturer’s procedure (Waters). Aliquots of each digest were TMT labeled. Label incorporation was verified to be >95% by LC-MS/MS and spectral counting. Samples were then pooled and lyophilized. The TMT channel layouts were: 126C: DMSO/Monoculture (Rep1); 127N: DMSO/ Coculture (Rep1); 127C: DMSO/Monoculture (Rep2); 128N: DMSO/Coculture (Rep2); 128C: DMSO/Monoculture (Rep3); 129N: DMSO/Coculture (Rep3); 129C: DMSO/Monoculture (Rep4); 130N: DMSO/Coculture (Rep4); 130C: Alectinib 1 μM/Monoculture (Rep1); 131N: Alectinib 1 μM/Coculture (Rep1); 131C: Alectinib 1 μM/Monoculture (Rep2); 132N: Alectinib 1 μM/Coculture (Rep2); 132C: Alectinib 1 μM/Monoculture (Rep3); 133N: Alectinib 1 μM/Coculture (Rep3); 133C: Alectinib 1 μM/Monoculture (Rep4); 134N: Alectinib 1 μM/Coculture (Rep4). After lyophilization, TMT-labeled peptides were redissolved in 200 μL of aqueous 20 mM ammonium formate, (pH 10.0). Basic pH reversed phase liquid chromatography (bRPLC) separation was performed on a XBridge 4.6 mm x 100 mm column packed with BEH C18 resin, 3.5 μm particle size, 130 Å pore size (Waters). bRPLC Solvent A was aqueous 2% acetonitrile with 5 mM ammonium formate (pH 10.0). Peptides were eluted by the following gradient: 5% Solvent B (aqueous 90% acetonitrile with 5 mM ammonium formate, pH 10.0) for 10 minutes, 5%- 15% B in 5 minutes, 15-40% B in 47 minutes, 40-100% B in 5 minutes and 100% B held for 10 minutes, followed by re-equilibration at 1% B. The flow rate was 0.6 mL/min, and 12 and 24 concatenated fractions were collected for phosphorylation and expression proteomics samples, respectively. Samples were dried by vacuum centrifugation (Thermo). Dried peptides were re-dissolved in immobilized metal affinity chromatography (IMAC) loading buffer containing aqueous 0.1% TFA and 85% acetonitrile. Phosphopeptides were enriched using IMAC resin (Cell Signaling Technology # 20432) washed with loading buffer on a Kingfisher (Thermo). Peptides were incubated with 5 μL of resin for 30 minutes at room temperature with gentle agitation. The IMAC resin was washed twice with loading buffer followed by wash buffer (aqueous 80% acetonitrile, 0.1% TFA). Phosphopeptides were eluted with aqueous 50% acetonitrile, 2.5% ammonia. The volume was reduced to 20 μl via vacuum centrifugation.

### LC-MS/MS conditions for proteomics analysis

A nanoflow ultra-high-performance liquid chromatograph and nano electrospray orbitrap mass spectrometer (RSLCnano and Q Exactive HF-X, Thermo) were used for LC-MS/MS. The sample was loaded onto a pre-column (C18 PepMap100, 100 μm ID x 2 cm length packed with C18 reversed-phase resin, 5 μm particle size, 100 Å pore size) and washed for 8 minutes with aqueous 2% acetonitrile and 0.1% formic acid. Trapped peptides were eluted onto the analytical column, (C18 PepMap100, 75 μm ID x 25 cm length, 2 μm particle size, 100 Å pore size, Thermo). A 120- minute gradient was programmed as: 95% solvent A (aqueous 2% acetonitrile + 0.1% formic acid) for 8 minutes, solvent B (aqueous 90% acetonitrile + 0.1% formic acid) from 5% to 38.5% in 90 minutes, then solvent B from 50% to 90% B in 7 minutes and held at 90% for 5 minutes, followed by solvent B from 90% to 5% in 1 minute and re-equilibration for 10 minutes using a flow rate of 300 nL/min. Spray voltage was 1900 V. Capillary temperature was 275°C. S lens RF level was set at 40. The top 20 tandem mass spectra were collected in a data-dependent manner. The resolution for MS and MS/MS were set at 60,000 and 45,000 respectively. Dynamic exclusion was 15 seconds for previously sampled peaks. MaxQuant (version 1.6.14.0) was used to identify peptides using the UniProt human database (March 2020) and quantify the TMT reporter ion intensities. Up to 2 missed trypsin cleavages were allowed. The mass tolerance was 20 ppm and 4.5 ppm for the first and the main search, respectively. Reporter ion mass tolerance was set to 0.003 Da, minimal precursor intensity fraction to 0.75. Carbamidomethyl cysteine was set as fixed modification. Phosphorylation on serine/threonine/tyrosine, deuterium-labeled lysine (+8) and methionine oxidation were set as variable modifications. Both peptide spectral match (PSM) and protein false discovery rates (FDR) were set at 0.05. Match between runs was activated to carry identifications across samples.

### Proteomics data normalization and analysis

For TMT-labeled total protein expression experiments (no injection replicates), each sample was normalized with iterative rank order normalization (IRON)^76^ (iron_generic-proteomics) against the median sample for its respective dataset (light: 128N, heavy: 130C). For TMT-labeled phosphorylation experiments, for each injection replicate, the first DMSO channel (126C) was used as the reference channel for IRON normalization. In order to remove systematic differences in signal between multiplex injection replicates, for each injection replicate, a computational pool of normalized log2 abundances was calculated for each row, then used to convert all abundances into log2 ratios vs. each row’s computational pool. Next, computational pools were averaged across injection replicates and used to scale the log2 ratios back into de-batched abundances.

Average log2 abundances were calculated from biological replicates after first averaging any technical replicates. Log2 ratios between conditions were calculated by subtracting average log2 abundances. Two-group p-values were calculated using two-sided, unequal-variance Welch’s T-tests from the individual averaged technical replicates within each condition. Differences between differences, “delta-delta” comparisons (used for comparing coculture to monoculture), were similarly calculated, appropriately propagating the standard error and degrees of freedom throughout the 4-group p-value calculations. For each comparison, standard deviations of the log2 ratios were calculated separately to be used as fold-change cutoffs during differential expression analysis. Comparisons were determined to be differentially expressed if all of the following criteria were met: row is assigned to at least one protein from the target species, |log2 ratio| > 2σ from the average value, and p-value < 0.05. Scores were calculated as the geometric mean of the |log2 ratio| and -log10(p-value), multiplied by the sign of the log2 ratio. Rows were sorted on |Score|, direction of change, and differential expression status. Volcano plots of expression and phospho-proteomics data were generated in GraphPad Prism. KSEA analysis was performed using the KSEA app (casecpb.shinyapps.io/ksea/).^41^ Pathway analyses were performed and plotted using gProfiler package within R studio (version 4.2.2).

### Flow cytometry

Nuclight Red H3122 cells in mono- or coculture with immortalized CAF12 were treated with DMSO or 1 μM alectinib for 3 hours and harvested. Cells were dissociated with Accumax (Fisher Scientific, #NC9870954), washed twice with PBS and fixed with 4% formaldehyde. Cells were then permeabilized with ice-cold 100% methanol or 0.1% Triton X-100 followed by rinsing and incubation with primary antibody phospho-ERK (Cell Signaling, #4370) or act-ITGB1 (Millipore Co, #MAB2079-AF647) for 1 hour respectively on ice. Cells for phospho-ERK determination were rinsed 3 times, incubated with Alexa Fluor488-conjugated secondary antibodies (Cell Signaling, #4412) for 45 min at room temperature in the dark and rinsed another two times with PBS. Cells were analyzed using a LSRII (BD Biosciences) Flow Cytometer. All analyses were performed using FlowJo (BD Biosciences). Singlet, live and red cell gating was performed consistently for each file. The geometric mean of the phosphorylation signal was used.

### RNA interference

H3122 cells were reverse transfected using Lipofectamine RNAiMAX Transfection Reagent (Thermo Fisher Scientific, no. 13778150) according to the manufacturer’s instructions. In brief, 20 nM final concentration siRNA (Horizon Discovery Bioscience, *ITGB1* ONTARGETplus SMARTpool, #L-004506-00-0005), or the control non-targeting pool (Horizon Discovery Bioscience, si-NT, #D-001810-10-20) were incubated for 20 min in a six-well dish containing 5 μL lipofectamine RNAiMAX in 500 μL of Opti-MEM (Gibco, #31985062) medium. Subsequently, 0.6e^6^ H3122 cells were added in 2 mL RPMI10. After 72 hours at 37°C and 5% CO2, cells were plated in a 384-well microtiter plate at 1,000 cells/well overnight. Then, drug or DMSO media were added for continuous treatment until 72 hours followed by red nuclei count via Incucyte as described earlier. In parallel, cells were collected for immunoblotting.

### Fibronectin-coating rescue experiment

For immunoblotting upon fibronectin (FN1) and drug treatment, 6-well plates were pre-coated with human purified FN1 (FisherScientific, #341635) at 2.5 μg/cm^2^ according to manufacturer’s instructions. First, 0.6e^6^ H3122 cells were suspended in 1.5 mL media and plated in uncoated or coated wells overnight. Then, media were aspirated followed by adding drug/media solution (1.5 mL). Cells were then harvested after 1 hour of drug treatment for immunoblotting analysis. For cell viability assays, 384-well plates were precoated and cell viability was read out by an Incucyte as described above.

### CRISPR-Cas9 targeting of *ITGB1*

Lentivirus particles carrying the Lenti_EFS-Cas9-P2A-HygR plasmid (Addgene, #164134) were generated using a 3^rd^ generation virus packaging system (ABM, LV053) by transfecting HEK293T cells. Targeted cells were transduced with Lenti_EFS-Cas9-P2A-HygR viral supernatant. The transduced cells were then split and selected with 50 μg/mL hygromycin for 7 days. Immunoblotting for the Flag tag was performed to confirm the expression of Cas9 in transduced cells. Cas9 expressing cells were then transduced with lenti-guide-DNA viral supernatant. The *ITGB1* forward and reverse small guide RNA sequences were: sgRNA1: Forward: 5’- CACCGTTTGTGCACCACCCACAATT-3’; Reverse: 5’-AAACAATTGTGGGTGGTGCACAAAC- 3’; sgRNA2: Forward: 5’-CACCGTGCTGTTCCTTTGCTACGGT-3’; Reverse: 5’- AAACACCGTAGCAAAGGAACAGCAC-3’.^77^ *AAVS1* targeting sgRNA was used as control: Forward: 5’- CACCGACTGTTGACGGCGGCGATGT-3’; Reverse: 5’-AAACACATCGCCGCCGTCAACAGTC-3’.^78^ Oligonucleotides were synthesized by Integrated DNA Technologies, Inc. and were phosphorylated and annealed before insertion into the BsmBI-digested lenti-CRISPR backbone LRG2.1 (Addgene, #108098). Lentiviruses carrying the LRG2.1-guide DNA construct were produced as described above and used for transduction in Cas9-expressing cells. Then, cells were imaged to confirm efficient viral transduction as indicated by green fluorescence. Pools of sg*ITGB1*-Cas9-Nuclight Red cancer cells and sg*AAVS1*-Cas9-Nuclight Red cancer cells were then sampled for immunoblotting to determine ITGB1 expression levels and subjected to cell viability assays as described above.

### Mouse allograft

Transgenic mouse *EML4-ALK*-positive cells (mEA1)^51^ were subcutaneously injected into 4- to 6- week-old Bl/6 recipient mice (Jackson Laboratory). Each animal received two contralateral injections containing 0.25 million mEA1 cells suspended in 100 μL of 1:1 mix of RPMI/BME type 3. Mice were randomized into treatment and control groups subjected to daily oral gavage or *i.p.* with vehicle (0.5% HPMC and 0.1% Tween80), capmatinib (3 mg/kg, MedChemExpress, #HY- 13404, oral), cilengitide (5 mg/kg, MedChemExpress, #HY-16141, i.p.), alectinib (20 mg/kg, Hoffmann-La Roche, oral), alectinib plus capmatinib, alectinib plus cilengitide or triple combination. Vehicle and single drug control groups had 3 mice each while combination treatment groups had 5 mice each. Tumor diameters, measured by electronic calipers, and animal weights were measured weekly. For tumor volume calculations, cubic shape of tumors was assumed, and relative volume were normalized to the volume at day 1 before drug treatment. After 4 weeks of treatment, animals were euthanized and tumors were weighed.^19^

### Immunohistochemistry

Formalin-fixed, paraffin embedded tumors were cut as 5 micron sections. Deparaffinized tissue slices were blocked in PBS with 10% goat serum for 30 min at room temperature, then incubated with phospho-S6 antibody for 1 hour (1:200, CST, #4858) followed by biotinylated anti-rabbit secondary antibody (1:100, Vector Labs, #BA-1000) for 30 min. Staining was developed using Vectastain ABC HRP Kit (Vector Labs, PK-6100), following manufacturer’s protocol. Cytoseal XYL mounting media (Epredia) was used to mount the slides. Images were scanned with Aperio ScanScope XT Slide Scanner (Leica). Segmentation was performed using an artificial intelligence based semi-automated deep learning quantification method (Aiforia release 5.6, Aiforia Inc, Cambridge, MA). For pS6 segmentation model, training set consisted of 35 H3122 images. Area fraction was used as a metric for pS6 content. Epithelial tissue was segmented from whole tissue excluding connective tissue, necrosis, and stroma and separated onto pS6+ (DAB-positive) and pS6- (DAB-negative) areas. pS6+ rate was calculated as a percentage of pS6+ area from total epithelial tissue.

### Correlation and survival analyses

Gene expression and survival data of NSCLC patients were fetched from ‘oncosg’ dataset from CBioPortal (www.cbioportal.com) on 2022/10/5. Data were filtered based on genetic alteration information to pool data of patients with *EGFR*/*ROS1*/*ALK* alterations only. Kaplan-Meier analysis was performed and plotted for patients with survival length of upper quartile vs lower quartile using R. The same dataset that was used for correlation analysis between fibroblast activation protein (FAP-α) expression level and tumor stages. FAP-α expression levels were compared between patients at stage I and II versus patients of stage III and IV. T test was performed to calculate p-values. Graph plotting and statistics were performed in R (version 4.3.0).

## Acknowledgements

We would like to thank Dr. N.P. Gauthier (Memorial Sloan-Kettering Cancer Center), Dr. K.-C. Han, Dr. O. Deng and Dr. D. Abate-Daga (all Moffitt Cancer Center) for advice on designing/optimizing CTAP experiments and reagents. Funding: This work was supported by the NIH/NCI R01 CA219347 (to U.R. and E.B.H.); the Florida Department of Health Bankhead-Coley Cancer Research Program, award no. 30-20450-9901 (to A.M.); Miles for Moffitt; Moffitt Innovative Core Technology Committee; and the H. Lee Moffitt Cancer Center & Research Institute. We further wish to acknowledge the Moffitt Lung Cancer Center of Excellence and the Moffitt Proteomics & Metabolomics, Analytic Microscopy, Molecular Genomics, and Flow Cytometry Core Facilities, as well as the Biostatistics and Bioinformatics Shared Resource, which are supported in part by the National Cancer Institute (award no. P30-CA076292) as a Cancer Center Support Grant. The Proteomics and Metabolomics Core is also supported by the Moffitt Foundation.

## Author contributions

Conception and design: Q.H., L.L.R.R., J.M.K., E.B.H., A.M., and U.R. Development of methodology: Q.H., L.L.R.R. and G.M.W. Acquisition of data: Q.H., L.L.R.R., B.D., X.L., B.F., and A.M. Analysis and interpretation of data: Q.H., L.L.R.R., B.D., D.M., E.A.W., and B.F. Administrative, technical, or material support: X.L., N.C., J.L.K., and R.C.D. Writing of the manuscript: Q.H. and U.R. Study supervision: U.R. All authors read, edited, and approved the final manuscript.

## Data and materials availability

The MS proteomics data have been deposited to the ProteomeXchange Consortium via the PRIDE^79^ partner repository with the dataset identifiers PXD053444 and 10.6019/PXD053444. All other data needed to evaluate the conclusions in the paper are present in the paper or the Supplementary Materials and are available from the corresponding author upon reasonable request.

## References

1 Bedard, P. L., Hyman, D. M., Davids, M. S. & Siu, L. L. Small molecules, big impact: 20 years of targeted therapy in oncology. Lancet 395, 1078–1088 (2020). 10.1016/S0140-6736(20)30164-1

2 Koivunen, J. P. et al. EML4-ALK fusion gene and efficacy of an ALK kinase inhibitor in lung cancer. Clin Cancer Res 14, 4275–4283 (2008). 10.1158/1078-0432.CCR-08-0168

3 Solomon, B., Varella-Garcia, M. & Camidge, D. R. ALK gene rearrangements: a new therapeutic target in a molecularly defined subset of non-small cell lung cancer. J Thorac Oncol 4, 1450–1454 (2009). 10.1097/JTO.0b013e3181c4dedb

4 Engelman, J. A. & Settleman, J. Acquired resistance to tyrosine kinase inhibitors during cancer therapy. Curr Opin Genet Dev 18, 73–79 (2008). 10.1016/j.gde.2008.01.004

5 Lovly, C. M. & Shaw, A. T. Molecular pathways: resistance to kinase inhibitors and implications for therapeutic strategies. Clin Cancer Res 20, 2249–2256 (2014). 10.1158/1078-0432.CCR-13-1610

6 Lin, J. J., Gainor, J. F., Lam, V. K. & Lovly, C. M. Unlocking the Next Frontier in Precision Oncology: Addressing Drug-Tolerant Residual Disease. Cancer Discov 14, 915–919 (2024). 10.1158/2159-8290.CD-24-0374

7 Hata, A. N. et al. Tumor cells can follow distinct evolutionary paths to become resistant to epidermal growth factor receptor inhibition. Nat Med 22, 262–269 (2016). 10.1038/nm.4040

8 Ramirez, M. et al. Diverse drug-resistance mechanisms can emerge from drug-tolerant cancer persister cells. Nat Commun 7, 10690 (2016). 10.1038/ncomms10690

9 Sharma, S. V. et al. A chromatin-mediated reversible drug-tolerant state in cancer cell subpopulations. Cell 141, 69–80 (2010). 10.1016/j.cell.2010.02.027

10 Bhowmick, N. A., Neilson, E. G. & Moses, H. L. Stromal fibroblasts in cancer initiation and progression. Nature 432, 332–337 (2004). 10.1038/nature03096

11 Ohlund, D., Elyada, E. & Tuveson, D. Fibroblast heterogeneity in the cancer wound. J Exp Med 211, 1503–1523 (2014). 10.1084/jem.20140692

12 Moghal, N. et al. Single-Cell Analysis Reveals Transcriptomic Features of Drug-Tolerant Persisters and Stromal Adaptation in a Patient-Derived EGFR-Mutated Lung Adenocarcinoma Xenograft Model. J Thorac Oncol 18, 499–515 (2023). 10.1016/j.jtho.2022.12.003

13 Biffi, G. & Tuveson, D. A. Diversity and Biology of Cancer-Associated Fibroblasts. Physiol Rev 101, 147–176 (2021). 10.1152/physrev.00048.2019

14 Chen, Y., McAndrews, K. M. & Kalluri, R. Clinical and therapeutic relevance of cancer-associated fibroblasts. Nat Rev Clin Oncol 18, 792–804 (2021). 10.1038/s41571-021-00546-5

15 McMillin, D. W., Negri, J. M. & Mitsiades, C. S. The role of tumour-stromal interactions in modifying drug response: challenges and opportunities. Nat Rev Drug Discov 12, 217–228 (2013). 10.1038/nrd3870

16 Harbinski, F. et al. Rescue screens with secreted proteins reveal compensatory potential of receptor tyrosine kinases in driving cancer growth. Cancer Discov 2, 948–959 (2012). 10.1158/2159-8290.CD-12-0237

17 Wilson, T. R. et al. Widespread potential for growth-factor-driven resistance to anticancer kinase inhibitors. Nature 487, 505–509 (2012). 10.1038/nature11249

18 Tape, C. J. et al. Oncogenic KRAS Regulates Tumor Cell Signaling via Stromal Reciprocation. Cell 165, 910–920 (2016). 10.1016/j.cell.2016.03.029

19 Remsing Rix, L. L., et al. IGF-binding proteins secreted by cancer-associated fibroblasts induce context-dependent drug sensitization of lung cancer cells. Sci Signal 15, eabj5879 (2022). 10.1126/scisignal.abj5879

20 Shaw, A. T. et al. Ceritinib in ALK-rearranged non-small-cell lung cancer. N Engl J Med 370, 1189–1197 (2014). 10.1056/NEJMoa1311107

21 Gadgeel, S. M. et al. Safety and activity of alectinib against systemic disease and brain metastases in patients with crizotinib-resistant ALK-rearranged non-small-cell lung cancer (AF-002JG): results from the dose-finding portion of a phase 1/2 study. Lancet Oncol 15, 1119–1128 (2014). 10.1016/S1470-2045(14)70362-6

22 Gettinger, S. N. et al. Activity and safety of brigatinib in ALK-rearranged non-small-cell lung cancer and other malignancies: a single-arm, open-label, phase 1/2 trial. Lancet Oncol 17, 1683–1696 (2016). 10.1016/S1470-2045(16)30392-8

23 Yao, Z. et al. TGF-beta IL-6 axis mediates selective and adaptive mechanisms of resistance to molecular targeted therapy in lung cancer. Proc Natl Acad of Sci U S A 107, 15535–15540 (2010). 10.1073/pnas.1009472107

24 Shintani, Y. et al. IL-6 Secreted from Cancer-Associated Fibroblasts Mediates Chemoresistance in NSCLC by Increasing Epithelial-Mesenchymal Transition Signaling. J Thorac Oncol 11, 1482–1492 (2016). 10.1016/j.jtho.2016.05.025

25 Patel, S. A. et al. IL6 Mediates Suppression of T- and NK-cell Function in EMT-associated TKI-resistant EGFR-mutant NSCLC. Clin Cancer Res 29, 1292–1304 (2023). 10.1158/1078-0432.CCR-22-3379

26 Alsterda, A. et al. Salubrinal Exposes Anticancer Properties in Inflammatory Breast Cancer Cells by Manipulating the Endoplasmic Reticulum Stress Pathway. Front Oncol 11, 654940 (2021). 10.3389/fonc.2021.654940

27 Lane, D., Matte, I., Rancourt, C. & Piche, A. Osteoprotegerin (OPG) protects ovarian cancer cells from TRAIL-induced apoptosis but does not contribute to malignant ascites-mediated attenuation of TRAIL-induced apoptosis. J Ovarian Res 5, 34 (2012). 10.1186/1757-2215-5-34

28 Straussman, R. et al. Tumour micro-environment elicits innate resistance to RAF inhibitors through HGF secretion. Nature 487, 500–504 (2012). 10.1038/nature11183

29 Yamada, T. et al. Paracrine receptor activation by microenvironment triggers bypass survival signals and ALK inhibitor resistance in EML4-ALK lung cancer cells. Clin Cancer Res 18, 3592–3602 (2012). 10.1158/1078-0432.CCR-11-2972

30 Yano, S. et al. Hepatocyte growth factor induces gefitinib resistance of lung adenocarcinoma with epidermal growth factor receptor-activating mutations. Cancer Res 68, 9479–9487 (2008). 10.1158/0008-5472.CAN-08-1643

31 Hu, H. et al. Three subtypes of lung cancer fibroblasts define distinct therapeutic paradigms. Cancer Cell 39, 1531–1547 e1510 (2021). 10.1016/j.ccell.2021.09.003

32 Hrustanovic, G. et al. RAS-MAPK dependence underlies a rational polytherapy strategy in EML4-ALK-positive lung cancer. Nat Med 21, 1038–1047 (2015). 10.1038/nm.3930

33 Gauthier, N. P. et al. Cell-selective labeling using amino acid precursors for proteomic studies of multicellular environments. Nat Methods 10, 768–773 (2013). 10.1038/nmeth.2529

34 Tape, C. J. et al. Cell-specific labeling enzymes for analysis of cell-cell communication in continuous co-culture. Mol Cell Proteomics 13, 1866–1876 (2014). 10.1074/mcp.O113.037119

35 Emdal, K. B. et al. Integrated proximal proteomics reveals IRS2 as a determinant of cell survival in ALK-driven neuroblastoma. Sci Signal 11 (2018). 10.1126/scisignal.aap9752

36 Zhang, G. et al. Coupling an EML4-ALK-centric interactome with RNA interference identifies sensitizers to ALK inhibitors. Sci Signal 9, rs12 (2016). 10.1126/scisignal.aaf5011

37 Voena, C. et al. The tyrosine phosphatase Shp2 interacts with NPM-ALK and regulates anaplastic lymphoma cell growth and migration. Cancer Res 67, 4278–4286 (2007). 10.1158/0008-5472.CAN-06-4350

38 Vaishnavi, A. et al. EGFR Mediates Responses to Small-Molecule Drugs Targeting Oncogenic Fusion Kinases. Cancer Res 77, 3551–3563 (2017). 10.1158/0008-5472.CAN-17-0109

39 Zhao, Y., Yang, Y., Xu, Y., Lu, S. & Jian, H. AZD 0530 sensitizes drug-resistant ALK- positive lung cancer cells by inhibiting SRC signaling. FEBS Open Bio 7, 472–476 (2017). 10.1002/2211-5463.12162

40 Katayama, R. et al. Mechanisms of acquired crizotinib resistance in ALK-rearranged lung Cancers. Sci Transl Med 4, 120ra117 (2012). 10.1126/scitranslmed.3003316

41 Wiredja, D. D., Koyuturk, M. & Chance, M. R. The KSEA App: a web-based tool for kinase activity inference from quantitative phosphoproteomics. Bioinformatics 33, 3489–3491 (2017). 10.1093/bioinformatics/btx415

42 Bernfield, M. & Sanderson, R. D. Syndecan, a developmentally regulated cell surface proteoglycan that binds extracellular matrix and growth factors. Philos Trans R Soc Lond B Biol Sci 327, 171–186 (1990). 10.1098/rstb.1990.0052

43 Evanko, S. P., Potter-Perigo, S., Petty, L. J., Workman, G. A. & Wight, T. N. Hyaluronan Controls the Deposition of Fibronectin and Collagen and Modulates TGF-beta1 Induction of Lung Myofibroblasts. Matrix Biol 42, 74–92 (2015). 10.1016/j.matbio.2014.12.001

44 Fujisaki, T. et al. CD44 stimulation induces integrin-mediated adhesion of colon cancer cell lines to endothelial cells by up-regulation of integrins and c-Met and activation of integrins. Cancer Res 59, 4427–4434 (1999).

45 McFarlane, S., McFarlane, C., Montgomery, N., Hill, A. & Waugh, D. J. CD44-mediated activation of alpha5beta1-integrin, cortactin and paxillin signaling underpins adhesion of basal-like breast cancer cells to endothelium and fibronectin-enriched matrices. Oncotarget 6, 36762–36773 (2015). 10.18632/oncotarget.5461

46 Bill, H. M. et al. Epidermal growth factor receptor-dependent regulation of integrin-mediated signaling and cell cycle entry in epithelial cells. Mol Cell Biol 24, 8586–8599 (2004). 10.1128/MCB.24.19.8586-8599.2004

47 Mitra, A. K. et al. Ligand-independent activation of c-Met by fibronectin and alpha(5)beta(1)-integrin regulates ovarian cancer invasion and metastasis. Oncogene 30, 1566–1576 (2011). 10.1038/onc.2010.532

48 Maroun, C. R., Naujokas, M. A., Holgado-Madruga, M., Wong, A. J. & Park, M. The tyrosine phosphatase SHP-2 is required for sustained activation of extracellular signal-regulated kinase and epithelial morphogenesis downstream from the met receptor tyrosine kinase. Mol Cell Biol 20, 8513–8525 (2000). 10.1128/MCB.20.22.8513-8525.2000

49 Hynes, R. O. Integrins: bidirectional, allosteric signaling machines. Cell 110, 673–687 (2002). 10.1016/s0092-8674(02)00971-6

50 Kapp, T. G. et al. A Comprehensive Evaluation of the Activity and Selectivity Profile of Ligands for RGD-binding Integrins. Sci Rep 7, 39805 (2017). 10.1038/srep39805

51 Maddalo, D. et al. In vivo engineering of oncogenic chromosomal rearrangements with the CRISPR/Cas9 system. Nature 516, 423–427 (2014). 10.1038/nature13902

52 Wang, W. et al. Crosstalk to stromal fibroblasts induces resistance of lung cancer to epidermal growth factor receptor tyrosine kinase inhibitors. Clin Cancer Res 15, 6630–6638 (2009). 10.1158/1078-0432.CCR-09-1001

53 Cords, L. et al. Cancer-associated fibroblast phenotypes are associated with patient outcome in non-small cell lung cancer. Cancer Cell 42, 396–412 e395 (2024). 10.1016/j.ccell.2023.12.021

54 Lambrechts, D. et al. Phenotype molding of stromal cells in the lung tumor microenvironment. Nat Med 24, 1277–1289 (2018). 10.1038/s41591-018-0096-5

55 Pellinen, T. et al. Fibroblast subsets in non-small cell lung cancer: Associations with survival, mutations, and immune features. J Natl Cancer Inst 115, 71–82 (2023). 10.1093/jnci/djac178

56 McMillin, D. W. et al. Tumor cell-specific bioluminescence platform to identify stroma-induced changes to anticancer drug activity. Nat Med 16, 483–489 (2010). 10.1038/nm.2112

57 Ishibashi, M. et al. CD200-positive cancer associated fibroblasts augment the sensitivity of Epidermal Growth Factor Receptor mutation-positive lung adenocarcinomas to EGFR Tyrosine kinase inhibitors. Sci Rep 7, 46662 (2017). 10.1038/srep46662

58 Hirata, E. et al. Intravital imaging reveals how BRAF inhibition generates drug-tolerant microenvironments with high integrin beta1/FAK signaling. Cancer Cell 27, 574–588 (2015). 10.1016/j.ccell.2015.03.008

59 Haderk, F. et al. Focal adhesion kinase-YAP signaling axis drives drug-tolerant persister cells and residual disease in lung cancer. Nat Commun 15, 3741 (2024). 10.1038/s41467-024-47423-0

60 Zhang, S. et al. The Potent ALK Inhibitor Brigatinib (AP26113) Overcomes Mechanisms of Resistance to First- and Second-Generation ALK Inhibitors in Preclinical Models. Clin Cancer Res 22, 5527–5538 (2016). 10.1158/1078-0432.CCR-16-0569

61 Zou, H. Y. et al. PF-06463922 is a potent and selective next-generation ROS1/ALK inhibitor capable of blocking crizotinib-resistant ROS1 mutations. Proc Natl Acad Sci U S A 112, 3493–3498 (2015). 10.1073/pnas.1420785112

62 Hu, Q. et al. Differential Chemoproteomics Reveals MARK2/3 as Cell Migration-Relevant Targets of the ALK Inhibitor Brigatinib. Chembiochem, e202200766 (2023). 10.1002/cbic.202200766

63 Liao, Y. et al. Differential network analysis of ROS1 inhibitors reveals lorlatinib polypharmacology through co-targeting PYK2. Cell Chem Biol 31, 284–297 e210 (2024). 10.1016/j.chembiol.2023.09.011

64 Kuenzi, B. M. et al. Polypharmacology-based ceritinib repurposing using integrated functional proteomics. Nat Chem Biol 13, 1222–1231 (2017). 10.1038/nchembio.2489

65 Cooper, J. & Giancotti, F. G. Integrin Signaling in Cancer: Mechanotransduction, Stemness, Epithelial Plasticity, and Therapeutic Resistance. Cancer Cell 35, 347–367 (2019). 10.1016/j.ccell.2019.01.007

66 Barrow-McGee, R. et al. Beta 1-integrin-c-Met cooperation reveals an inside-in survival signalling on autophagy-related endomembranes. Nat Commun 7, 11942 (2016). 10.1038/ncomms11942

67 Jahangiri, A. et al. Cross-activating c-Met/beta1 integrin complex drives metastasis and invasive resistance in cancer. Proc Natl Acad Sci U S A 114, E8685–E8694 (2017). 10.1073/pnas.1701821114

68 Palve, V., Liao, Y., Remsing Rix, L. L. & Rix, U. Turning liabilities into opportunities: Off-target based drug repurposing in cancer. Semin Cancer Biol 68, 209–229 (2021). 10.1016/j.semcancer.2020.02.003

69 Noh, K. W. et al. Integrin beta3 Inhibition Enhances the Antitumor Activity of ALK Inhibitor in ALK-Rearranged NSCLC. Clin Cancer Res 24, 4162–4174 (2018). 10.1158/1078-0432.CCR-17-3492

70 Bokhari, A. A. et al. Novel human-derived EML4-ALK fusion cell lines identify ribonucleotide reductase RRM2 as a target of activated ALK in NSCLC. Lung Cancer 171, 103–114 (2022). 10.1016/j.lungcan.2022.07.010

71 Mediavilla-Varela, M., Boateng, K., Noyes, D. & Antonia, S. J. The anti-fibrotic agent pirfenidone synergizes with cisplatin in killing tumor cells and cancer-associated fibroblasts. BMC Cancer 16, 176 (2016). 10.1186/s12885-016-2162-z

72 Campeau, E. et al. A versatile viral system for expression and depletion of proteins in mammalian cells. PLoS One 4, e6529 (2009). 10.1371/journal.pone.0006529

73 Park, J. et al. Recording of elapsed time and temporal information about biological events using Cas9. Cell 184, 1047–1063 e1023 (2021). 10.1016/j.cell.2021.01.014

74 Tarumoto, Y. et al. LKB1, Salt-Inducible Kinases, and MEF2C Are Linked Dependencies in Acute Myeloid Leukemia. Mol Cell 69, 1017-1027 e1016 (2018). 10.1016/j.molcel.2018.02.011

75 Hayer, A. et al. Engulfed cadherin fingers are polarized junctional structures between collectively migrating endothelial cells. Nat Cell Biol 18, 1311–1323 (2016). 10.1038/ncb3438

76 Welsh, E. A., Eschrich, S. A., Berglund, A. E. & Fenstermacher, D. A. Iterative rank-order normalization of gene expression microarray data. BMC Bioinformatics 14, 153 (2013). 10.1186/1471-2105-14-153

77 Cheng, S. L., X.; Yuan, Y.; Jia, C.; Chen, L.; Gao, Q.; Lu, Z.; Yang, R.; Nie, G.; Yang, J.; Duan, W.; Xiao, L.; Hou, Y. ITGB1 Enhances the Proliferation, Survival, and Motility in Gastric Cancer Cells. Microscopy and Microanalysis 27, 1192–1201 (2021). 10.1017/S1431927621012393

78 Lin, A. et al. Off-target toxicity is a common mechanism of action of cancer drugs undergoing clinical trials. Sci Transl Med 11 (2019). 10.1126/scitranslmed.aaw8412

79 Perez-Riverol, Y. et al. The PRIDE database resources in 2022: a hub for mass spectrometry-based proteomics evidences. Nucleic Acids Res 50, D543–D552 (2022). 10.1093/nar/gkab1038

