## Supplementary information for "Cancer-associated fibroblasts confer ALK inhibitor resistance in *EML4-ALK*-driven lung cancer via concurrent integrin and MET signaling"

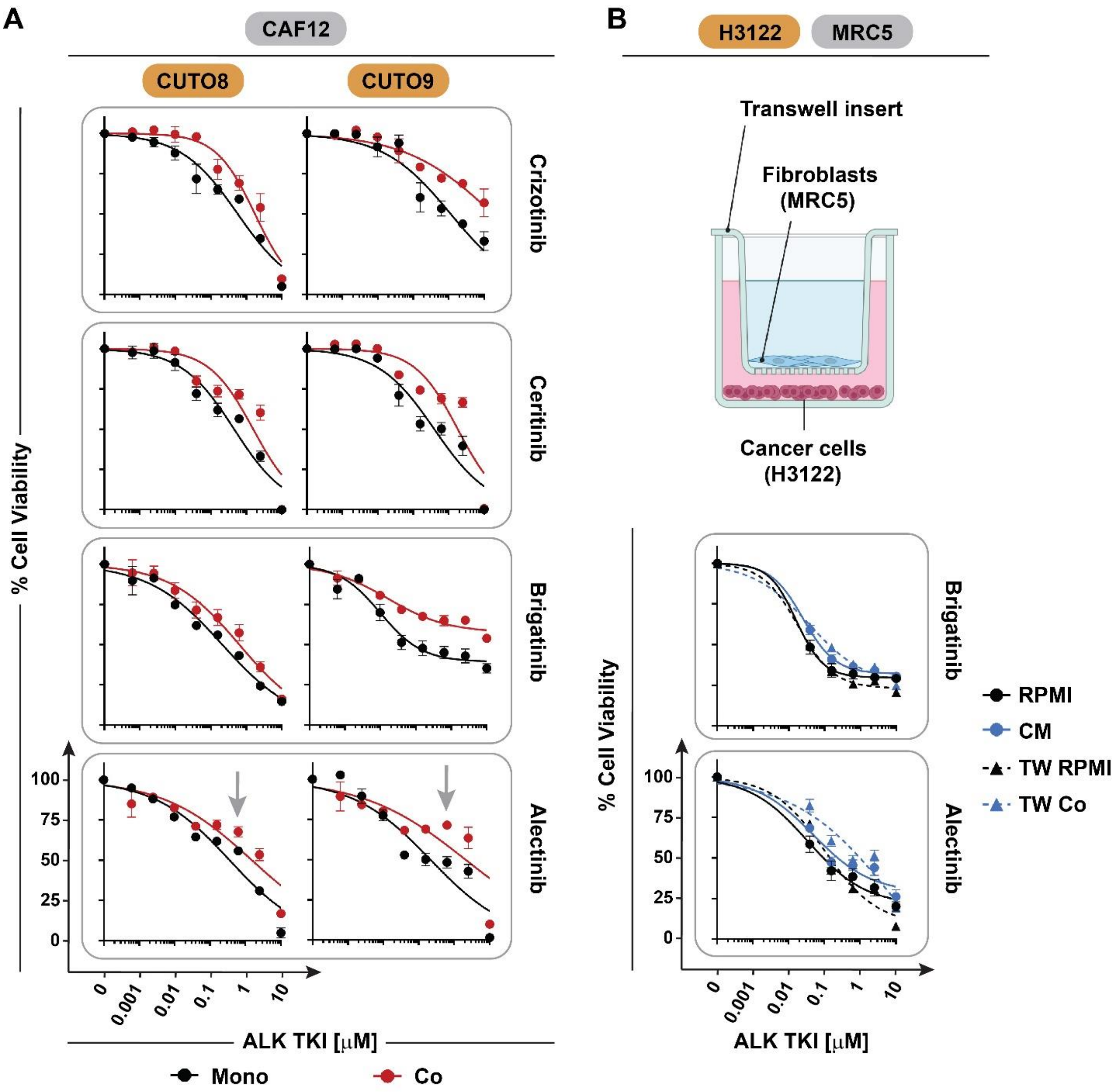


**Figure S1. Drug sensitivity of *EML4-ALK*-positive NSCLC cells in physical and transwell coculture with fibroblasts. A.** Viability of CUTO8 and CUTO9 *EML4-ALK*+ NSCLC cells after 72 hours of treatment with different ALK TKIs in CAF12 coculture. Grey arrows indicate the alectinib concentration, which was selected for statistical analysis using one-way ANOVA w/ Holm-Sidak's multiple comparison correction (see **Figure 1F**). n = 3. **B.** Viability of H3122 cells in MRC5 CM or transwell (TW) coculture after 72 hours of treatment with brigatinib or alectinib at indicated concentrations.

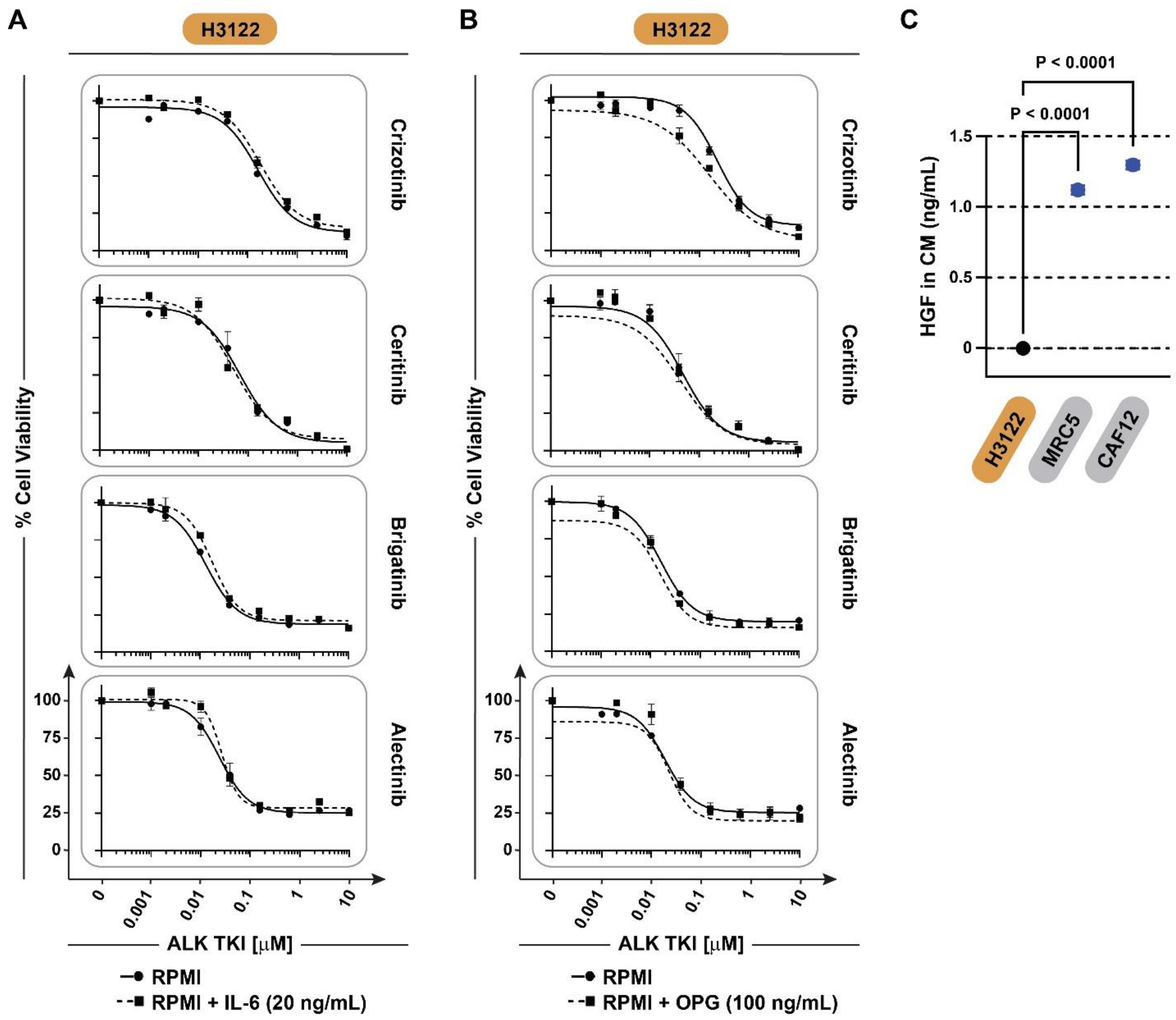


**Figure S2. Effects of fibroblast-secreted factors on *EML4-ALK*-positive NSCLC cells. A./B.** Viability of H3122 cells upon treatment with ALK TKIs at indicated concentrations and IL-6 (20 ng/mL; **A**) or OPG (100 ng/mL; **B**) for 72 hours. n = 2. **C.** Quantification of HGF concentration in CM from H3122, MRC5 and CAF12 determined by ELISA. *P* values were determined using one-way ANOVA w/ Holm-Sidak's multiple comparison correction. n = 3.

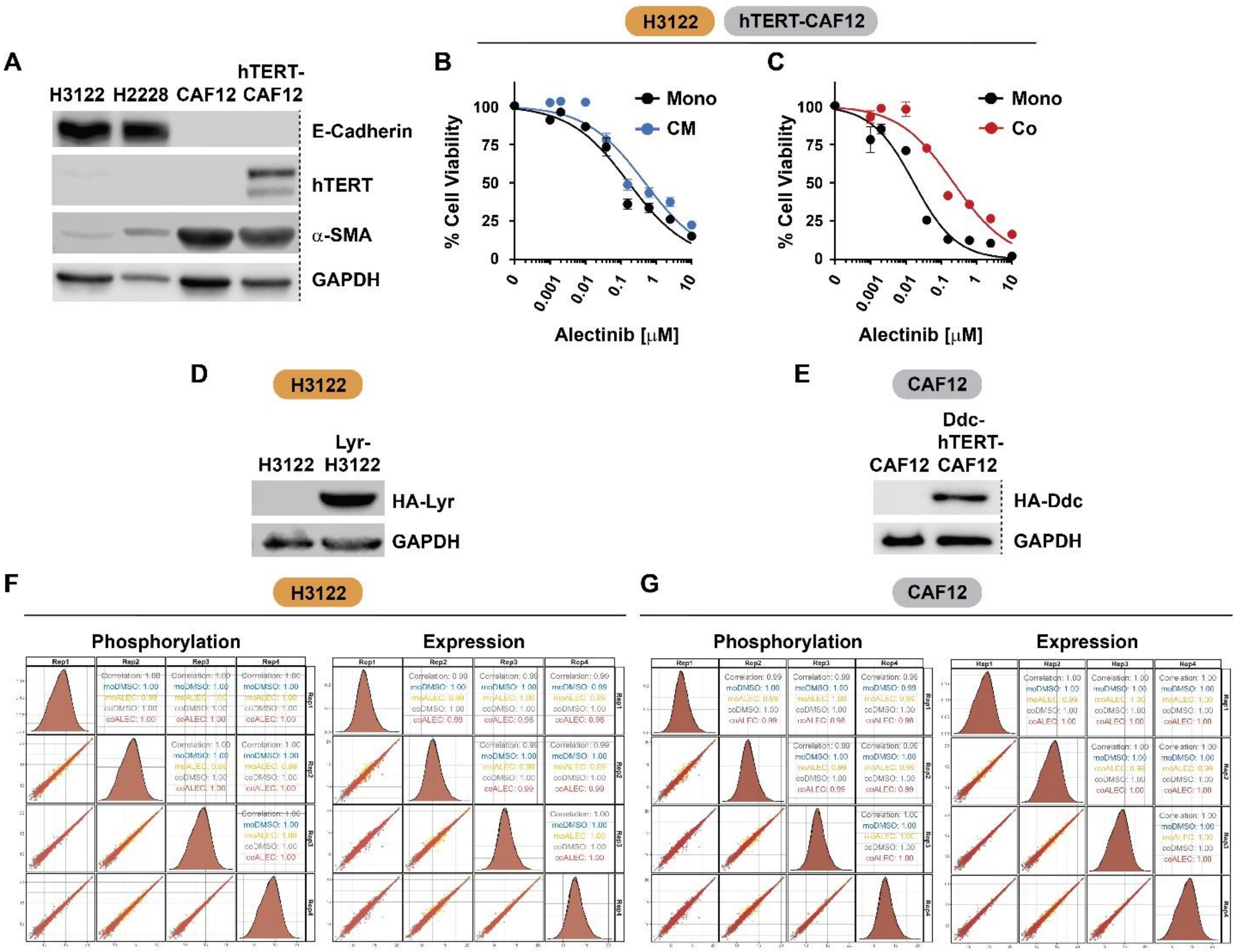


**Figure S3. CTAP cell engineering and data reproducibility. A.** Immunoblot analysis of hTERT, E-cadherin, and α-SMA in immortalized CAF12 cells compared to parental CAF12 and NSCLC cells. n = 3. **B.** Viability of H3122 cells in hTERT-CAF12 CM upon treatment with alectinib at indicated concentrations for 72 hours as determined by CellTiter-Glo viability assay. n = 3. **C.** Viability of H3122 cells in hTERT-CAF12 coculture upon treatment with alectinib at indicated concentrations for 72 hours as determined by live-cell imaging. n = 2. **D./E.** Immunoblot analysis of HA-Lyr and HA-Ddc expression in H3122 (**D**) and immortalized CAF12 (**E**), respectively, compared to their parental cells. **F./G.** Correlation plots across replicates of phospho- and expression proteomics in H3122 (**F**) and CAF12 (**G**).

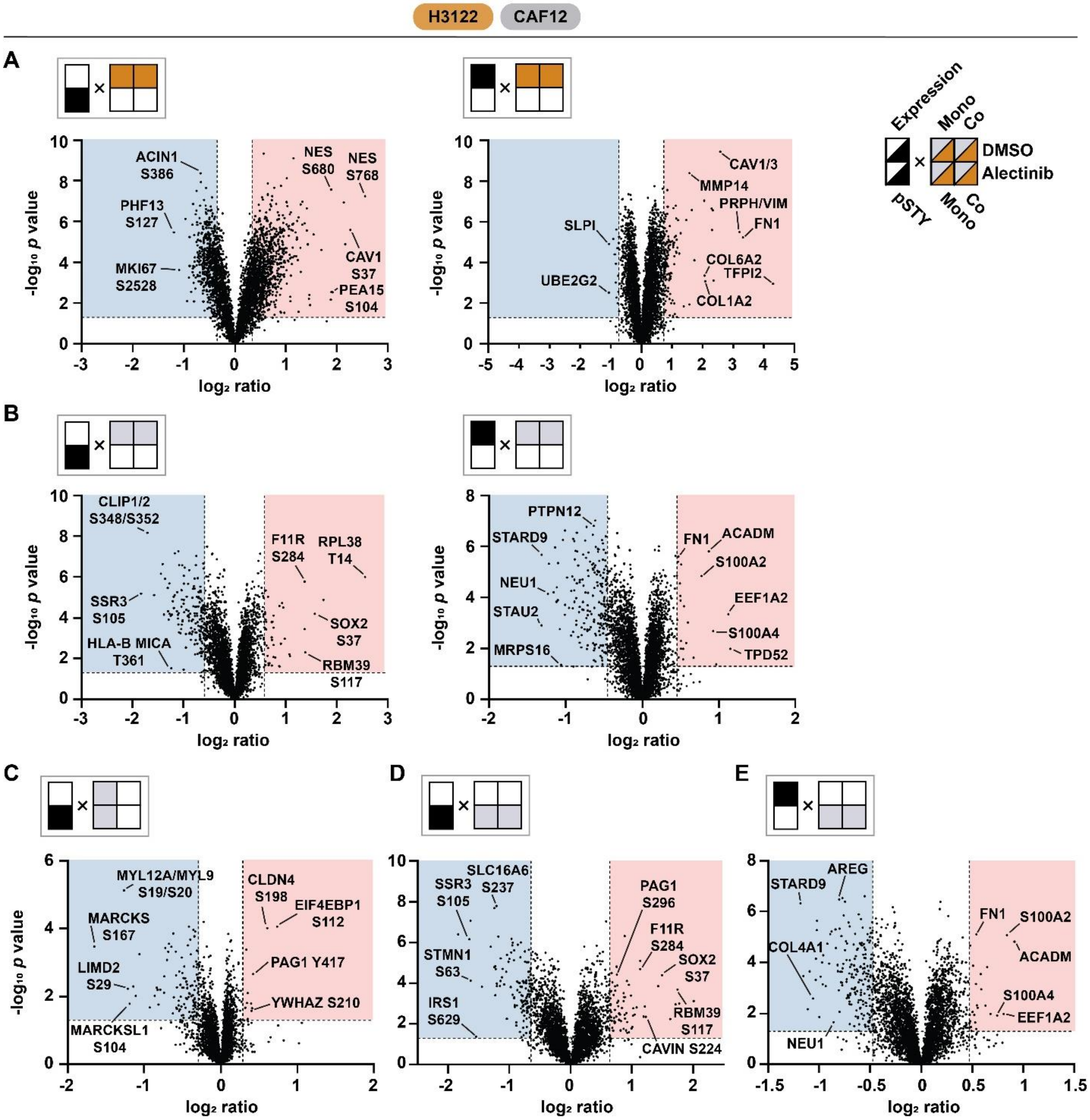


**Figure S4. Coculture-based phospho- and expression proteomics. A.** Global phosphorylation (left) and protein expression (right) changes in H3122 cells upon treatment with DMSO in mono- compared to CAF12 coculture. **B.** Global phosphorylation (left) and protein expression (right) changes in CAF12 cells upon treatment with DMSO in mono- compared to H3122 coculture. **C.** Global phosphorylation changes in CAF12 cells upon treatment with alectinib compared to DMSO in monoculture. **D.** Global phosphorylation changes in CAF12 cells in mono- compared to H3122 coculture upon treatment with alectinib. **E.** Protein expression changes in CAF12 cells in mono- compared to H3122 coculture upon treatment with alectinib. **A.-E.** Significantly regulated signals are indicated by highlighted areas (up: red; down: blue).

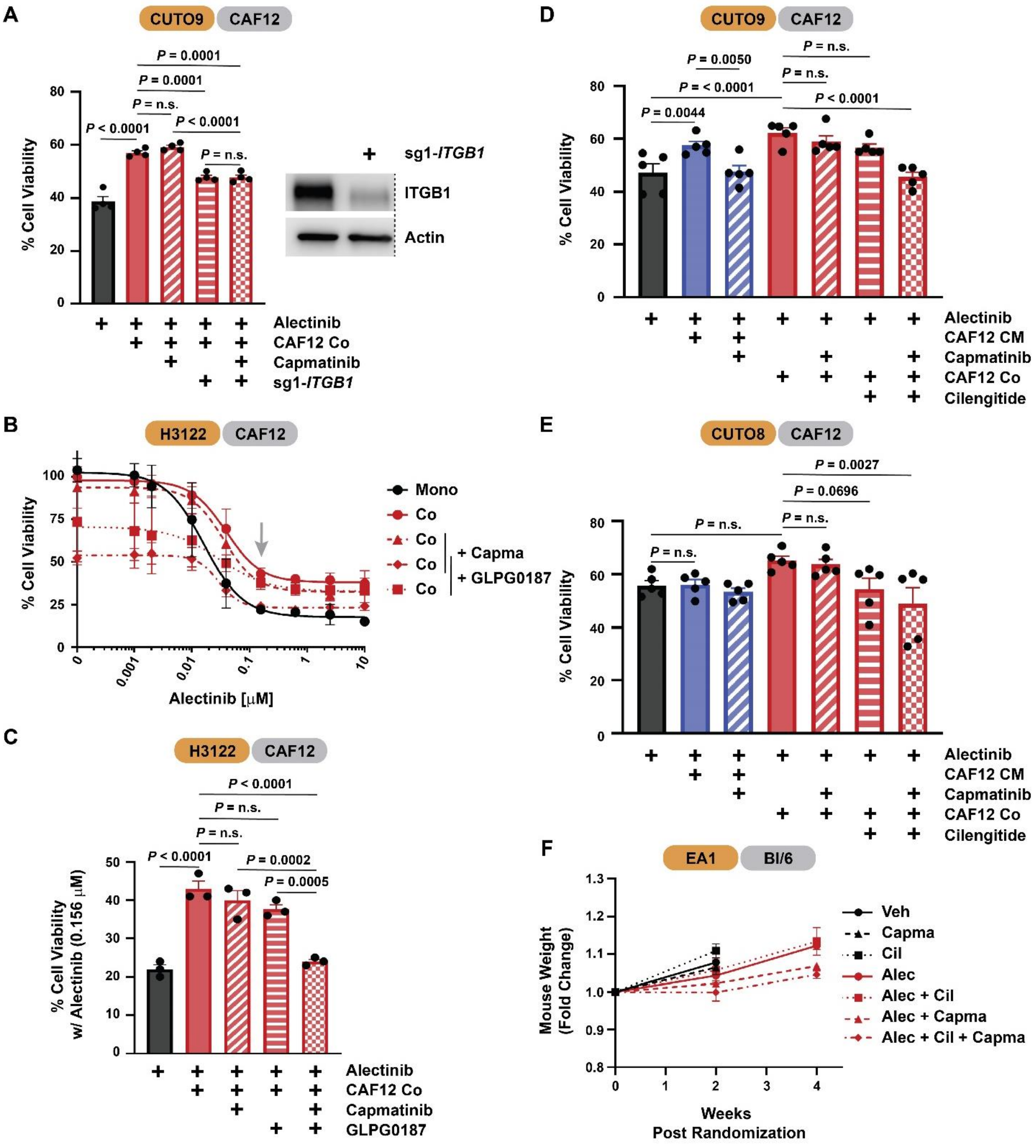


**Figure S5. Functional effects of dual targeting of ITGB1 and MET. A.** Viability of CUTO9 cells upon CRISPR-mediated knockout of *ITGB1* vs. *AAVS1* control and treatment with alectinib (2.5 μM) and capmatinib (0.2 μM) in mono- or CAF12 coculture. *P* values were determined using one-way ANOVA w/ Holm-Sidak's multiple comparison correction. n = 4. **B./C.** Viability of H3122 cells upon treatment with alectinib, capmatinib (0.2 μM) and the integrin inhibitor GLPG0187 (1 nM) in mono- or CAF12 coculture. Grey arrow indicates alectinib concentration selected for statistical analysis using one-way ANOVA w/ Holm-Sidak's multiple comparison correction (**C**). n = 3. **D./E.** Viability of CUTO9 (**D**) and CUTO8 (**E**) cells upon treatment with alectinib (2.5 μM), capmatinib (0.2 μM) and cilengitide (1 μM) in mono- or

CAF12 coculture. *P* values were determined using one-way ANOVA w/ Holm-Sidak's multiple comparison correction. n = 5. **F.** Relative weekly mouse weight post randomization of allograft model upon drug treatment (see **Figure 6A**).

### Other Supplementary Data

**Data file S1. Global CTAP-based phosphoproteomics data in H3122 cells.** Multiplex channel layout of samples and analysis of the phosphoproteomics data for heavy CTAP-labeled H3122 cells.

**Data file S2. Global CTAP-based phosphoproteomics data in CAF12 cells.** Multiplex channel layout of samples and analysis of the phosphoproteomics data for light CTAP-labeled hTERT-CAF12 cells.

**Data file S3. Global CTAP-based expression proteomics data in H3122 cells.** Multiplex channel layout of samples and analysis of the expression proteomics data for heavy CTAP-labeled H3122 cells.

**Data file S4. Global CTAP-based expression proteomics data in CAF12 cells.** Multiplex channel layout of samples and analysis of the expression proteomics data for light CTAP-labeled hTERT-CAF12 cells.
